# E4BP4 Safeguards Brown Fat Mitochondria from Obesity-Induced Fragmentation via Ceramide Repression

**DOI:** 10.1101/2025.05.19.652826

**Authors:** Fernando Valdivieso-Rivera, Vanessa O. Furino, Carlos E. Leher, Ariane M. Zanesco, Monara Kaélle Cruz, Flavia C. Gan, Adriana Leandra Santoro, Lara Regina-Ferreira, Giovanna Leite Santos, Tiago Gonçalves, Luiz Osório Leiria, Pedro M. Moraes-Vieira, Roger Frigério Castilho, Shingo Kajimura, Marcelo A. Mori, Licio A. Velloso, Carlos H. Sponton

**Affiliations:** Obesity and Comorbidities Research Center, Universidade Estadual de Campinas (UNICAMP), Campinas, Brazil; Graduate Program in Molecular and Morphofunctional Biology, Universidade Estadual de Campinas (UNICAMP), Campinas, Brazil; Laboratory of Immunometabolism, Department of Genetics, Evolution, Microbiology and Immunology-Institute of Biology, Universidade Estadual de Campinas (UNICAMP), Campinas, Brazil; Department of Biochemistry and Tissue Biology, Institute of Biology, Universidade Estadual de Campinas (UNICAMP), Campinas, Brazil; Department of Pathology, Universidade Estadual de Campinas (UNICAMP), Campinas, Brazil; Center for Research in Inflammatory Diseases (CRID), Ribeirão Preto Medical School, Universidade de São Paulo (FMRP-USP), Ribeirão Preto, Brazil; Experimental Medicine Research Cluster, Universidade Estadual de Campinas, Campinas (UNICAMP), Campinas, Brazil; Division of Endocrinology, Diabetes and Metabolism, Beth Israel Deaconess Medical Center and Harvard Medical School, and Howard Hughes Medical Institute, Boston, MA, USA; Department of Physiology, Ribeirão Preto Medical School, Universidade de São Paulo (FMRP-USP), Ribeirão Preto, Brazil

**Keywords:** Obesity, Brown Adipose Tissue, Mitochondrial Dynamics, Transcription Factors, Ceramides

## Abstract

Brown adipose tissue (BAT) counteracts obesity-related metabolic dysfunction through both thermogenic and non-thermogenic means. However, substantial evidence indicates that obesity negatively affects BAT mitochondrial morphology and oxidative capacity, impairing systemic energy homeostasis. Motivated by this apparent contradiction, we investigated the relationship between obesity and mitochondrial dynamics, as the underlying mechanisms remain incompletely understood. Here, we identified E4BP4 as a transcriptional repressor that prevents obesity-induced mitochondrial fragmentation and oxidative dysfunction by inhibiting ceramide synthesis in brown fat. Specifically, E4BP4 interacts with PRDM16 to repress Cers6 mRNA expression and consequently reduces C16:0 ceramide levels by binding to a 65 kb upstream enhancer region of the Cers6 gene. Notably, the preservation of mitochondrial integrity in BAT by E4BP4 gain-of-function improves systemic glucose homeostasis, independent of weight loss. Collectively, our findings establish E4BP4 as a molecular safeguard against obesity-induced mitochondrial fragmentation and oxidative dysfunction, primarily by suppressing ceramide synthesis in brown fat.

## INTRODUCTION

BAT is a metabolically active adipose depot characterized by high mitochondrial density and the capacity to dissipate energy as heat via uncoupling protein 1 (UCP1)-dependent and UCP1-independent mechanisms^1,2^. Numerous rodent studies have demonstrated that BAT-mediated non-shivering thermogenesis enhances whole-body energy expenditure, promotes weight loss, and counteracts obesity-related metabolic diseases^3^. However, whether BAT-induced thermogenesis significantly improves human systemic energy homeostasis remains debatable^4^. Emerging evidence suggests that BAT may influence systemic glucose homeostasis in non-thermogenic ways^3^. For example, BAT facilitates the catabolism of branched-chain amino acids (BCAAs), supplying nitrogen for hepatic biosynthesis of essential amino acids and glutathione^5^. Impaired BCAAs flux and catabolism within BAT mitochondria have been associated with insulin resistance independent of changes in energy expenditure or body weight^5^. Regardless of whether it is through non-shivering thermogenesis or other means, maintaining mitochondrial health is crucial for BAT’s beneficial effects on systemic energy homeostasis.

Given the central role of mitochondrial integrity in BAT function, understanding how mitochondrial morphology is regulated under metabolic stress is essential. Changes in mitochondrial architecture through fusion and fission events, collectively termed mitochondrial dynamics, represent adaptive responses to an organism’s energy supply and demand, directly influencing mitochondrial oxidative capacity^6^. Studies have indicated that obesity leads to an imbalance in mitochondrial dynamics, favoring fission events that result in mitochondrial fragmentation and reduced oxygen consumption in the liver^7^, muscle^8^, and in white adipose tissue (WAT)^9^. Mechanistically, recent evidence has demonstrated that obesity induces the expression and activity of RalA, a GTPase family protein, in WAT, driving Drp1-mediated mitochondrial fragmentation^9^. Ablation of RalA in WAT prevents weight gain and enhances fatty acid oxidation in diet-induced obese (DIO) mice. Interestingly, BAT mitochondrial morphology was unaffected by RalA activity, suggesting distinct regulatory mechanisms of mitochondrial morphology related to brown fat.

One potential upstream regulator of mitochondrial morphology during obesity is the accumulation of ceramides, which have emerged as key lipids in metabolic disease. Ceramides are bioactive sphingolipids that accumulate in various tissues during obesity, contributing significantly to cardiometabolic diseases, including type 2 diabetes, hepatic steatosis, and atherosclerosis^10–17^. Numerous pharmacological and genetic interventions in rodents have demonstrated that inhibiting *de novo* ceramide synthesis in adipose tissue improves systemic energy homeostasis^16,18–22^. Furthermore, ceramide accumulation is sufficient and necessary to disrupt BAT thermogenic function and cause metabolic dysfunction in DIO mice^19^.

Among various ceramide species, C16:0 ceramide has garnered particular attention due to its direct impact on mitochondrial morphology and oxidative metabolism. C16:0 ceramide buildup within BAT is associated with altered mitochondrial morphology and decreased mitochondrial respiration^19^. C16:0 ceramide synthesis is mediated by the enzymatic activity of ceramide synthase (Cers) isoforms 5 and 6^23^. In a large human cohort study, visceral WAT expression of Cers6 was positively correlated with higher body mass index (BMI), fat accumulation, and hyperglycemia^16^. Moreover, conditional deletion of Cers6, specifically in UCP1+ cells, enhances BAT thermogenic capacity and improves systemic energy homeostasis in DIO mice^16^. Several distinct mechanisms have been proposed, through which C16:0 ceramide influences mitochondrial morphology and oxidative capacity^17^. For instance, C16:0 ceramide directly interacts with mitochondrial fission factor (Mff), thus promoting Drp1-dependent mitochondrial fragmentation in hepatocytes^7^. Notably, hepatic deficiency of CerS6, but not CerS5, confers protection against weight gain, hepatic lipid accumulation, insulin resistance, and impaired mitochondrial oxidative function in DIO mice^7^.

Despite substantial evidence linking obesity-induced ceramide accumulation to mitochondrial damage in BAT, the underlying mechanism remains unclear. In this study, we identified E4 promoter-binding protein 4 (E4BP4), a member of the proline– and acid-rich (PAR) family of transcription factors, as a critical safeguarding factor against obesity-induced mitochondrial fragmentation, thereby preserving the mitochondrial oxidative capacity. E4BP4 is co-regulated by PRDM16 and directly represses Cers6 expression, leading to decreased C16:0 ceramide levels. Notably, ceramide repression by the E4BP4-PRDM16 transcriptional complex significantly improved systemic glucose homeostasis independent of body weight loss. These findings enhance our understanding of brown fat biology and shed light on the molecular mechanisms underlying obesity-induced mitochondrial damage.

## RESULTS

### E4BP4 is a transcription factor repressed in BAT of diet-induced obesity (DIO) mice

Although considerable advances have elucidated the roles of various transcription factors (TF) in the differentiation of brown and beige adipocytes as well as in the activation of thermogenic program, our understanding of TF functions in metabolic disorders remains unclear. Thus, to identify TF candidates that may be relevant in obesity, we analyzed public bulk RNA-seq datasets from interscapular BAT (iBAT) of chow-fed and DIO mice^24^. We investigated 289 curated TF (**Table S1**) previously identified in iBAT, using mass spectrometry^25^. Among the top ten downregulated TF in DIO versus chow diet mice, several were previously linked to brown and beige adipocyte development and the activation of thermogenic genes, including Nra4a1, Arntl1 (Bmal1), Zbtb16, and Ebf1^26–29^ (**Figure 1A**). E4bp4 caught our attention because it is a basic leucine zipper (bZIP) transcription factor that contains a unique repressor domain^30^ (**Figure 1B**). Moreover, single nucleotide polymorphisms of E4bp4 were positive associated (HuGE scores) with a higher body mass index (8.59 –log10 p-value), waist-to-hip ratio (6.39 –log10 p-value), and weight (12.86 –log10 p-value), based on GWAS studies derived from the Type 2 Diabetes Knowledge Portal (https://t2d.hugeamp.org/) (**Figure 1C**).

**Figure 1.**
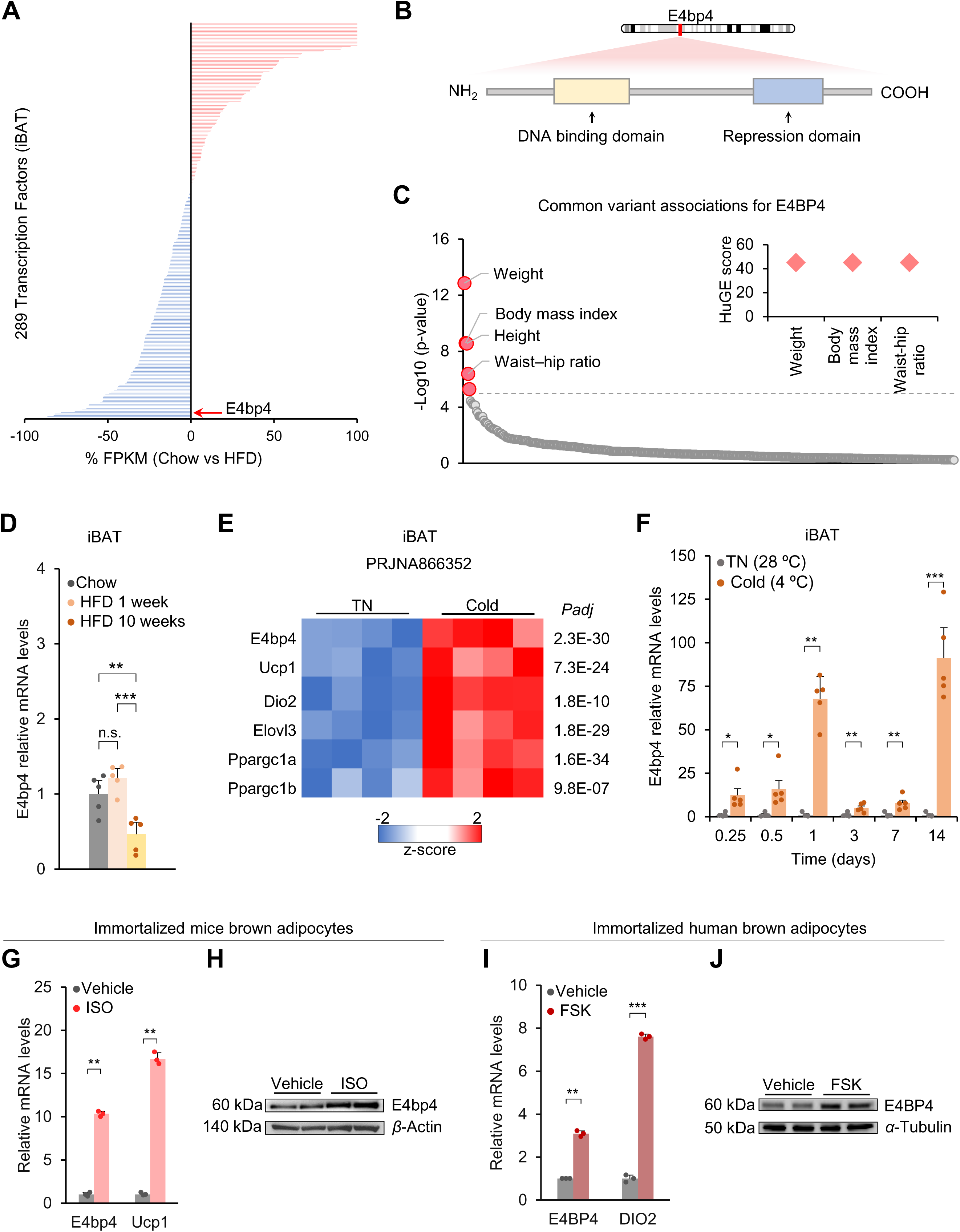
E4bp4 is a DNA-binding TF repressed by diet-induced obesity (DIO) and stimulated by cold and adrenergic signaling. A. FPKM-based quantification of expression changes (percentage) of 289 transcription factors in interscapular brown adipose tissue (iBAT) from mice fed a high-fat diet (HFD) versus a chow diet. Data are available in the Gene Expression Omnibus under the accession code GSE175608. B. Schematic representation of the transcription factor E4bp4. The basic leucine zipper (bZIP) domain and minimal repressor domain are indicated, illustrating regions critical for DNA binding and transcriptional repression. C. Left: The region page near the E4BP4 gene shows significant associations (above dash line) generated by bottom-line meta-analysis across all datasets in the Common Metabolic Diseases Knowledge Portal (CMDKP). Right: scores from the Human Genetic Evidence (HuGE) calculator on the gene page. D. Relative mRNA levels of E4pb4 in iBAT from HFD (1 and 10 weeks) and chow diet (10 weeks) control mice. n = 5 per group. E. *Heat map* of RNA-seq transcriptome of iBAT in animals under cold or thermoneutral (TN) stress. n = 4 per group. Data are represented as Z scores. Data are available from the BioProject: PRJNA866352. F. Relative mRNA levels of E4pb4 in iBAT from animals exposed to cold stress and thermoneutrality. n = 5 per group. G. Relative mRNA levels of E4pb4 and Ucp1 in brown adipocytes treated with isoproterenol (1 µM) or vehicle for 24 h. n = 3 per group. H. Western blotting for E4bp4 in brown adipocytes treated with isoproterenol (1 µM) or vehicle for 24 h. Representative results are shown from two independent experiments. Gel source data are presented in the Supplementary data. I. Relative mRNA levels of E4BP4 and DIO2 in immortalized human brown adipocytes treated with forskolin (20 µM) or vehicle for 24 h. n = 3 per group. J. Western blotting for E4BP4 in immortalized human brown adipocytes treated with forskolin (20 µM) or vehicle for 24 h. Representative results are shown from two independent experiments. Gel source data are presented in the Supplementary data. For D, F, G, and I, data are presented as the mean±SEM. Two-sided P values were calculated using unpaired Student’s t-tests (E, F, G, and I). Two-way ANOVA with Tukey’s multiple comparison test (D). *P < 0.05, **P < 0.01, ***P < 0.001.

E4BP4, also known as nuclear factor interleukin 3 regulated (Nfil3), belongs to the PAR subfamily of bZIP proteins, which includes DBP, HLF, and TEF^31^. It was originally described as a repressor that binds to a consensus sequence in the adenoviral E4 promoter^32^. Additionally, E4bp4 is an established component of the mammalian circadian clock, and prior findings suggest it may affect energy balance and metabolism^33,34^. However, its role in BAT remains unknown. To determine whether E4bp4 is suppressed by high-fat feeding, mice were fed an HFD for either one or ten weeks. After one week, E4bp4 mRNA levels remained unchanged, but after ten weeks, they were 54% reduced compared to the control (**Figure 1D**).

### E4BP4 expression is induced by cold stimulus and adrenergic signaling in brown fat

To investigate the regulation of E4bp4 in brown fat, we first analyzed publicly available iBAT bulk RNA-seq datasets^35^ from mice exposed to acute cold for 8h. The analysis revealed that E4bp4 expression, along with thermogenic genes (Ucp1, Dio2, Elovl3, Ppargc1α and Ppargc1β) were induced in mice under cold versus thermoneutrality (TN) (**Figure 1E, Table S2**). Alternatively, we performed gene expression analysis of iBAT in mice acutely (6h and 12h) or chronically (1-14 days) exposed to cold stress (4°C). As a control, mice were kept under TN. Our data revealed an oscillatory pattern of E4bp4 expression across different time points in mice under cold exposure (**Figure 1F**). A previous study similarly showed that E4bp4 expression rises early (at 2h), then declines in magnitude – while still remaining significantly related to control – during beige adipocyte differentiation^36^. Despite these temporal oscillations, our results confirmed that E4bp4 mRNA levels were higher in iBAT of mice under both acute and chronic cold exposure than in the control at all measured time points (**Figure 1F**).

To determine whether E4bp4 expression is mediated by β-adrenergic receptor (βAR) signaling, we treated immortalized brown adipocytes with the non-selective βAR agonist isoproterenol (ISO) for 24h. In this context, previous study has suggested that E4BP4 may play a role in cAMP-induced beige adipocyte differentiation^36^. Our results revealed an increase in E4bp4 mRNA levels (**Figure 1G**) and protein content (**Figure 1H**) in ISO compared to vehicle-treated cells. We also examined the regulation of E4bp4 expression in human brown adipocytes. The cells were treated with forskolin, an adenylate cyclase activator. Forskolin-treated human brown adipocytes had higher E4BP4 mRNA levels (**Figure 1I**) and protein content (**Figure 1J**) than those in the control group. Taken together, these data reveal that E4BP4 is induced in brown adipocytes by both cold stress and βAR signaling.

### E4BP4 overexpression (OE) prevented HFD-induced metabolic dysfunction

Next, we induced E4bp4 overexpression (E4bp4-OE) by injecting an adeno-associated vector (AAV) into the iBAT. AAV-GFP was used as the control. Mice were fed a HFD for 10 weeks at TN (28-29°C). The rationale for using this model was to provide housing conditions that closely resemble those of humans with minimal or no cold-induced thermogenesis^37^. Moreover, transgene expression by local AAV administration into brown fat is an alternative strategy to circumvent issues recently reported in the Ucp1 Cre mouse lineage, such as genetic abnormalities^38^ and non-adipose tissue transgene expression^39,40^. iBAT temperature was recorded using a thermal probe (IPTT300) throughout the 10 weeks of HFD feeding (**Figure S1A**). Our data revealed no differences in body weight (**Figure S1B**) or food intake (**Figure S1C**) between the two groups. Nonetheless, the iBAT temperature remained consistently higher in the E4BP4-OE group throughout the HFD feeding period (**Figure S1D**). AAV E4bp4 administration increased E4bp4 mRNA levels 63 fold compared to that in the control group (**Figure S1E**).

Next, we investigated the effects of E4bp4 gain-of-function on brown fat mass and morphology. iBAT mass was lower in the E4BP4 group than in the control group (**Figure S1F**). Histological analysis using hematoxylin and eosin (H&E) staining revealed a white adipocyte-like phenotype characterized by unilocular and large adipocytes in the control group. Conversely, E4BP4 displayed a typical multilocular morphology, with smaller adipocytes (**Figure S1G**). Moreover, immunofluorescence assay of an outer mitochondrial membrane protein (TOM20) revealed higher mitochondrial content in E4BP4 than in the control (**Figure S1H**).

Studies have demonstrated that E4BP4 is crucial for controlling the development and activity of immune cells^41^. Therefore, to avoid E4BP4 indirect effects on resident immune cells within iBAT, we next induced E4bp4 transgene expression using AAV under the control of the adiponectin promoter. In the first cohort, mice were fed an HFD for two weeks at TN (28-29°C), followed by indirect calorimetry (**Figure 2A**). This short-term HFD protocol minimizes the extensive iBAT remodeling (*whitening)* commonly observed in longer DIO studies^42,43^. We found increased energy expenditure (EE) in E4BP4-OE (**Figure 2B**), with no differences in body weight (**Figure S2A**) or tissue mass (**Figure S2B**). Moreover, the oxygen consumption rate (OCR) of iBAT was higher in the E4BP4-OE group (**Figure 2C**). E4bp4 mRNA was upregulated 46-fold relative to the control (**Figure 2D**).

**Figure 2.**
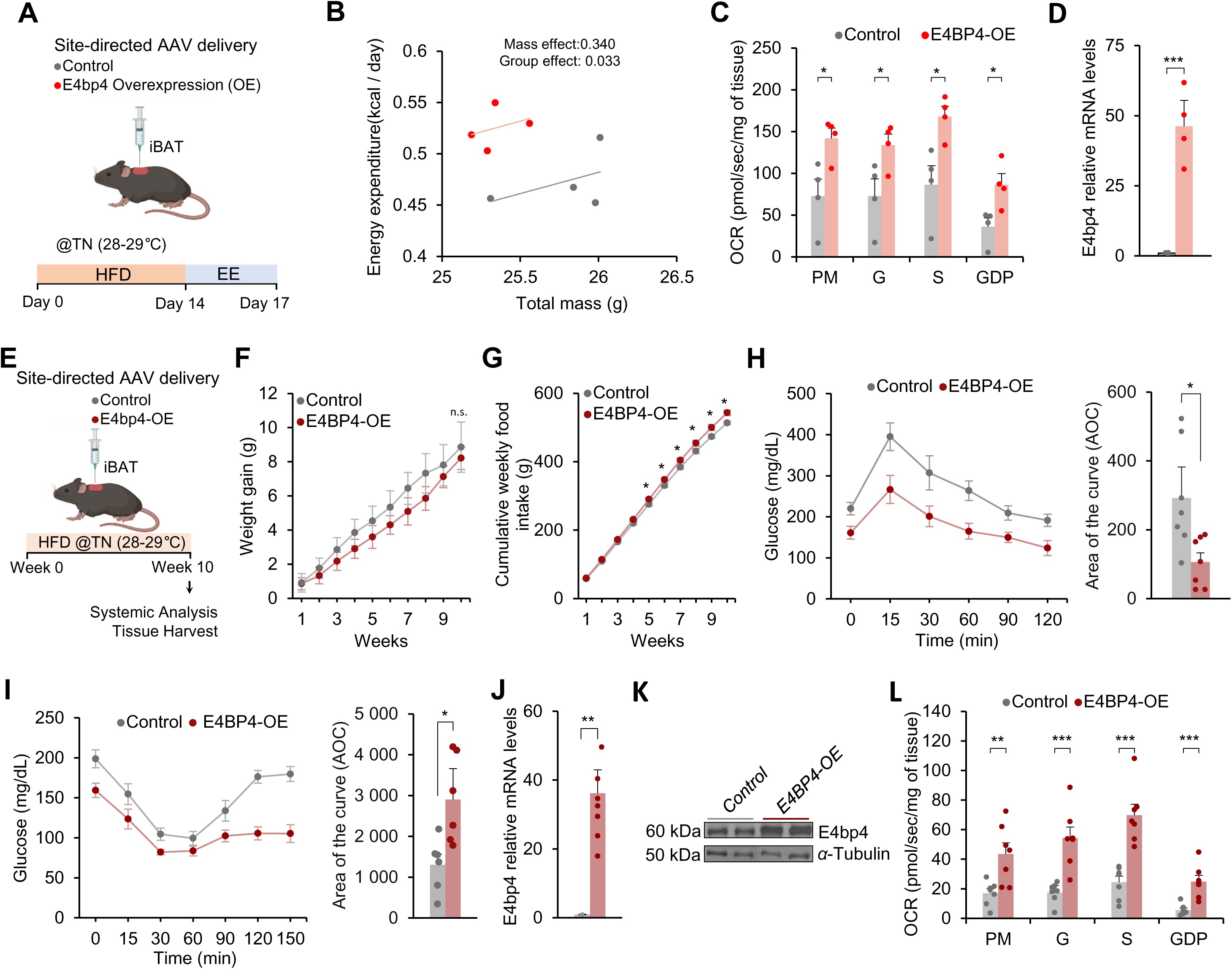
E4bp4 prevents obesity-induced impairment of systemic energy homeostasis. A. Schematic illustration of experimental design (A-D) for overexpression of E4bp4 (E4BP4-OE) or control in iBAT. The cartoon was created using a Biorender. B. Regression-based analysis of energy expenditure (EE) against body mass. Data were analyzed using a CaIR2-ANCOVA with EE as a dependent variable and body mass as a covariate. n = 4 per group. C. Mitochondrial oxygen consumption of iBAT in the presence of pyruvate/malate (PM), glutamate (G), succinate (S), and GDP-sensitive (UCP1 inhibitor). n = 4 per group. D. Relative mRNA levels of E4pb4 in iBAT from Control and E4BP4-OE after 2 weeks of HFD. n = 4 per group. E. Schematic illustration of the experimental design for overexpression of E4bp4 (E4BP4-OE) or Control in IBAT (F-L). The cartoon was created using a Biorender. F. Weight gain of Control and E4BP4-OE mice fed a HFD at 28-29°C. n = 7 per group. G. Cumulative weekly food intake of Control and E4BP4-OE mice fed a HFD at 28-29°C. n = 7 per group. H. Left: Glucose tolerance test of mice after 10 weeks of HFD. Right: Area of curve. n = 7 per group. I. Left: Insulin-tolerance test of mice after 10 weeks of HFD feeding. Right: Area of curve. n = 7 per group. J. Relative mRNA levels of E4pb4 in iBAT of Control and E4BP4-OE mice fed a HFD at 28-29°C. n = 7 per group. K. Western blotting for E4bp4 in iBAT of Control and E4BP4-OE mice fed an HFD at 28-29°C. Representative results are shown from two independent experiments. Gel source data are presented in the Supplementary data. L. Mitochondrial oxygen consumption of iBAT in the presence of pyruvate/malate (PM), glutamate (G), succinate (S), and GDP-sensitive (UCP1 inhibitor). n = 7 per group. For C, D, F, G, H, I, J, and L, data are presented as the mean±SEM. Two-sided P values were calculated using unpaired Student’s t-tests (C, D, H-L). Two-way ANOVA with Tukey’s multiple comparison test (F, G). *P < 0.05, **P < 0.01, ***P < 0.001.

In the second cohort, mice were exposed to a longer 10 weeks of HFD under TN (28-29°C) (**Figure 2E**). Consistent with our earlier findings (**Figure S1B**), body weight did not differ between the groups (**Figure 2F**). Interestingly, our data revealed a slight but significant increase in food intake in the E4BP4 group (**Figure 2G**). To further investigate the effects of the E4BP4 gain-of-function on glucose homeostasis, we performed glucose (GTT) and insulin (ITT) tolerance tests in both groups after 10 weeks of HFD feeding. Statistical analysis were performed using the area of the curve (AOC) as previously described^44^. Our data demonstrated a significant improvement in glucose tolerance (**Figure 2H**) and insulin sensitivity (**Figure 2I**) in E4BP4 compared with the control.

No differences were observed in subcutaneous inguinal adipose tissue (iWAT), liver, muscle, or heart mass between the two groups. In contrast, we found lower iBAT and gonadal adipose tissue (gWAT) mass for E4BP4 than for the control (**Figure S2C**). We observed increased iBAT E4bp4 mRNA levels (**Figure 2J**) and protein content (**Figure 2K**). It is worth noting that no E4bp4 transgene expression was detected in iWAT, gWAT, liver, or muscle (gastrocnemius) (**Figure S2D).** Similar to our earlier observations (**Figure S1G**), H&E staining of iBAT from E4BP4 exhibited a multilocular morphology with smaller adipocytes than those in the control (**Figure S2E**). Finally, the iBAT OCR remained significantly higher in E4BP4-OE than in the control (**Figure 2L**). Collectively, our data demonstrate that E4BP4 gain-of-function improves systemic energy homeostasis that is associated with changes in BAT mitochondrial content and oxidative capacity.

### E4BP4 prevents obesity-induced mitochondria fragmentation in brown fat

To understand how E4BP4 gain-of-function sustains higher brown fat mitochondrial content in DIO mice, we first analyzed publicly available single-cell RNA-sequencing datasets of iBAT. Our data revealed that the expression of E4bp4 was enriched in the brown adipocyte subpopulation along with Ucp1 and other genes related to the thermogenic program (**Figure S2F**). Therefore, we investigated whether E4bp4-OE induces mitochondrial biogenesis. However, we observed no differences in the mRNA levels of thermogenic or mitochondrial biogenesis genes such as Ucp1, Prdm16, Tfam, Nrf1, or Ppargc1α between the two groups (**Figure S2G**).

Obesity leads to an imbalance in mitochondrial dynamics, favoring fission events that result in fragmented mitochondria^9^. Fragmented and depolarized mitochondria activates mitophagy, leading to reduced mitochondria mass^6^. Thus, we explored whether E4BP4 gain-of-function prevents HFD-induced mitochondrial fragmentation in iBAT. Samples from the DIO mice, as described in Figure 2E, were subjected to transmission electron microscopy analysis. Our data demonstrated the appearance of smaller spherical mitochondria in the control group, which suggests more fragmented mitochondria^7,9^. In contrast, E4bp4 gain-of-function led to a higher frequency of elongated mitochondria (**Figures 3A – 3C**). Next, we investigated whether E4BP4 controls mitochondrial morphology in a *cell-autonomous* manner. E4bp4-OE was induced via AAV transduction, with AAV-GFP serving as the control in primary brown adipocytes (**Figure S3A**). As expected, E4bp4-OE led to a higher E4bp4 protein content (**Figure S3B**), with no effects on brown adipogenesis, compared to the control (**Figure S3C**). Lipid overload was induced *in vitro* by pretreating cells with palmitate (500 µM) for 24h and performing immunofluorescence analysis (**Figure 3D**). Confocal imaging revealed fewer fragmented mitochondria in E4bp4-OE cells than in the control (**Figure 3E**). Quantitative analysis confirmed that E4bp4-OE preserved mitochondrial volume and surface area, reduced sphericity, and maintained a higher number of branches and branch junctions, consistent with a more interconnected mitochondrial network (**Figure S4A**). Notably, no changes in the mitochondrial-to-nuclear DNA ratio were observed (**Figure S3D**) between the groups, which is consistent with our earlier data (**Figure S2G),** suggesting no induction of mitochondrial biogenesis by E4bp4 in DIO mice.

**Figure 3.**
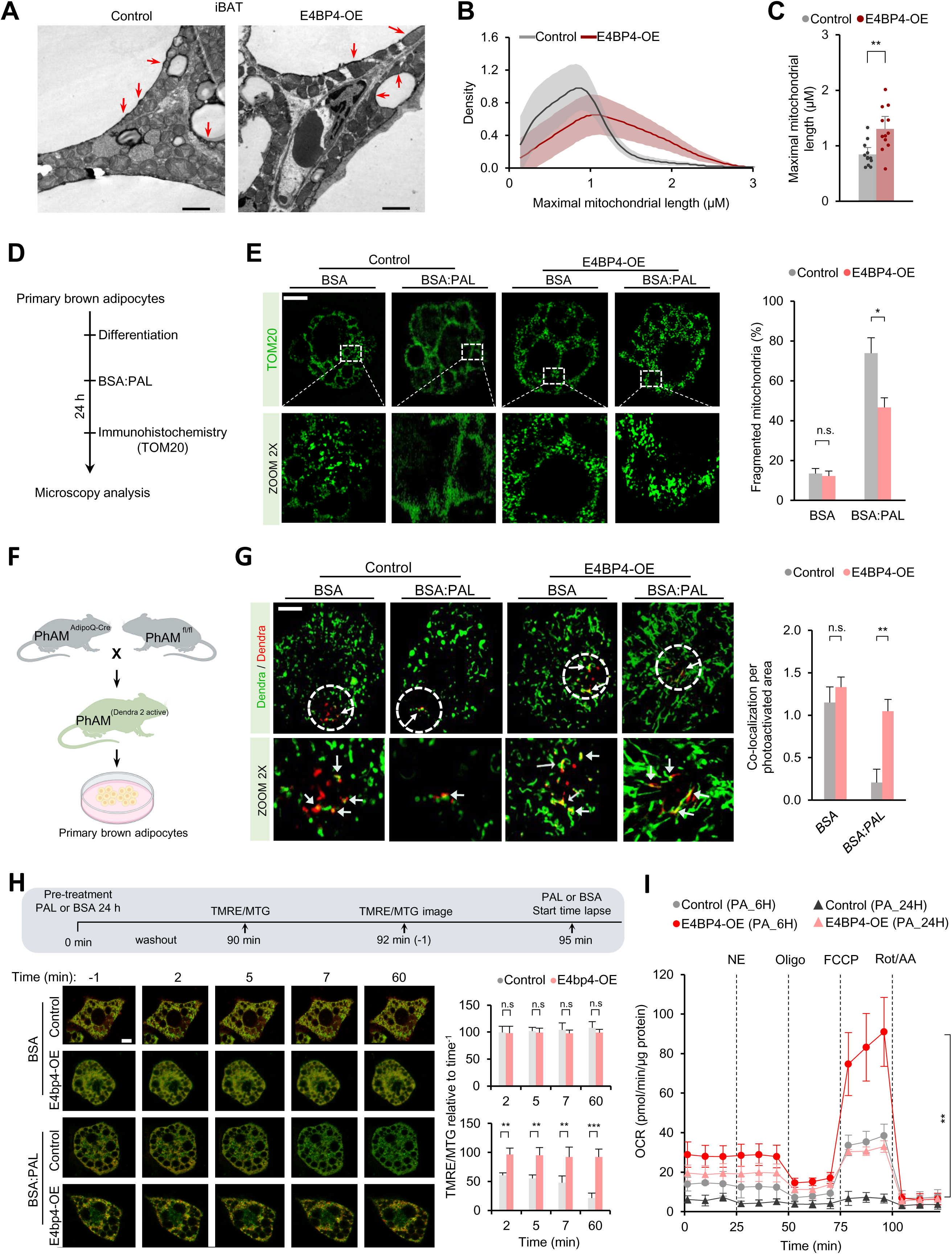
E4BP4 prevents obesity-induced mitochondria fragmentation. A. Transmission electron microscopy (TEM) of mitochondrial morphology in iBAT at 28-29°C in control and E4BP4-OE mice after 10 weeks of HFD feeding. Representative image. Scale bar: 2 µm. B. Histogram of maximal mitochondrial length of A. C. Quantification of maximal mitochondrial length (n = 3 per group). D. Schematic illustration of experimental design for E. The cartoon was created using a Biorender. E. Right: Images of mitochondrial morphology in primary brown adipocytes by TOM20 immunocytochemistry after 24 h of control treatment (BSA) or palmitate stimulation (BSA:PAL). Representative image. Scale bar, 10 µm. Left: Quantification of cells with a fragmented mitochondrial network (n = 3; 100-220 mitochondria from 10 images/group/experiment). F. Breeding strategy for PhAM^(Dendra2 active)^ mice and littermate control mice. Primary brown adipocytes were used in the experiments. The cartoon was created using a Biorender. G. Left: live-cell fluorescence of MitoDendra2 in primary brown adipocytes infected with AAV-control or E4BP4. Arrows show the colocalization of the active Dendra. Representative image. Scale bar, 12 µm. Right: Quantification of colocalization of Dendra active and not-activated. H. Top: Schematic illustration of the experimental design. Bottom left: Wave-like mitochondrial depolarization in primary brown adipocytes infected with AAV-Control and E4bp4 stained with TMRE (red) and MTG (green) stimulated with palmitate. Time-lapse imaging was performed at the onset of mitochondrial depolarization. Scale bar 5 µm. Bottom right: Quantification of depolarization occurring pre– and post-mitochondrial fragmentation (relative to minute –1). I. OCR in differentiated primary brown adipocytes expressing Control or E4bp4 after pretreatment for 6 or 24 h with control treatment (BSA) or palmitate stimulation (BSA:PAL). OCR values were normalized to total protein (µg). n = 5 per group. Norepinephrine (NE), oligomycin (oligo), carbonyl cyanide-p-trifluoromethoxyphenylhydrazone (FCCP), rotenone, and antimycin A (Rot/AA). For B, C, E, G, H, and I, data are presented as mean±SEM. Two-sided P-values were calculated using unpaired Student’s t-tests (C and H). Two-way ANOVA with Tukey’s multiple comparison test (E, G, and I). *P < 0.05, **P < 0.01, ***P < 0.001.

Alternatively, to investigate mitochondrial dynamics, we also used brown adipocytes derived from a previously established photo-activatable mitochondria (PhAM) reporter mouse model. PhAMfl/fl mice were crossed with adiponectin-Cre (AdipoQ-Cre) mice to drive conditional expression of the fluorescent protein Dendra2 specifically in adipocytes (**Figure 3F**). Dendra2 fluoresces green under baseline conditions but irreversibly switches to red fluorescence upon activation with a 405 nm laser. This enables tracking of mitochondrial fusion events by assessing the co-localization of pre-existing (green) and newly photo-activated (red) mitochondria. The iBAT was collected from these mice, and the stromal vascular fraction (SVF) was isolated and differentiated into brown adipocytes, which retained green-fluorescent mitochondria. Cells were then pre-treated with palmitate (500 μM) for 24 hours, followed by time-lapse microscopy (**Figure 3F**). Our results showed that palmitate treatment induced mitochondrial fragmentation in brown adipocytes, an effect that was prevented by E4BP4-OE (**Figure 3G; Video S1**). Notably, in vehicle-treated cells, we observed no increase in colocalized mitochondria in E4BP4 cells compared with the control. These findings suggest that the higher mitochondrial colocalization in the E4BP4 group is a consequence of fewer fission events than of higher fusion events.

We also evaluated whether the E4bp4 gain-of-function affects mitochondrial membrane potential (ΔΨm) of brown adipocytes. To this end, we used a combination of MitoTracker Green (MTG) and tetramethylrhodamine ethyl ester (TMRE) as previously reported^45^. Primary brown adipocytes were pretreated with palmitate (500 µM) for 24 h. After a washout period, we co-stained brown adipocytes with MTG and TMRE and reintroduced the palmitate-containing medium, followed by time-lapse microscopy (**Figure 3H**). Our findings revealed that the control cells experienced a noticeable decrease in ΔΨm within 2 min. In contrast, cells overexpressing E4bp4 maintained their ΔΨm for up to one hour (**Figure 3H**). Finally, we explored whether E4bp4 sustained ΔΨm could be translated into preserved mitochondrial oxidative function by performing an OCR analysis. Following pretreatment with palmitate (6h or 24h), we observed a modest but significant increase in both basal and norepinephrine-stimulated OCR in E4bp4-overexpressing cells compared with the control. Notably, E4bp4-OE elicited a robust increase in the maximal OCR induced by FCCP (**Figure 3I**). It is worth noting that a substantial proportional change in maximal OCR, exceeding that observed for other parameters, is likely attributable to enhanced mitochondrial substrate oxidation capacity, with alterations in mitochondrial uncoupling as a secondary consequence^46^. Overall, our data demonstrated that E4bp4 gain-of-function preserved brown fat ΔΨm and oxidative capacity by preventing mitochondrial fragmentation induced by lipid overload.

### E4BP4 OE suppress ceramide synthesis in brown fat

To investigate the underlying mechanisms, we conducted bulk RNA sequencing (RNA-seq) on iBAT samples obtained from the experiments shown in **Figure S1A**. Our differential gene expression analysis demonstrated that 722 and 217 genes were upregulated and downregulated, respectively (**Table S3**) in E4BP4 compared with those in the control. Enrichment pathway analysis (KEEG) revealed that “sphingolipids” was the most significantly downregulated pathway (**Figure 4A**). These findings have caught our attention because sphingolipids, particularly ceramides, have been shown to induce cellular and systemic metabolic dysfunction, by affecting mitochondria oxidative capacity^47^. Heatmap analysis demonstrated that the genes related to *de novo* synthesis of ceramides as well as “recycling pathways” were the most repressed genes in E4BP4 compared to the control (**Figure 4B**). To further investigate whether the decreased expression of ceramide biosynthetic genes might result in decreased sphingolipid levels, we performed mass spectrometry analysis of the “sphingolipidome” of iBAT from the experiments described in **Figure 2E**. Our data demonstrated that E4bp4 OE led to lower levels of C16:0 ceramide with no changes in other ceramide molecules (**Figure 4C**). Interestingly, we also observed an increase in the levels of other sphingolipid molecules, such as hexosylceramide (HexCer), sphingomyelin (SM), and lactosylceramide (LacCer), in E4BP4 compared to those in the control (**Figure 4C**).

**Figure 4.**
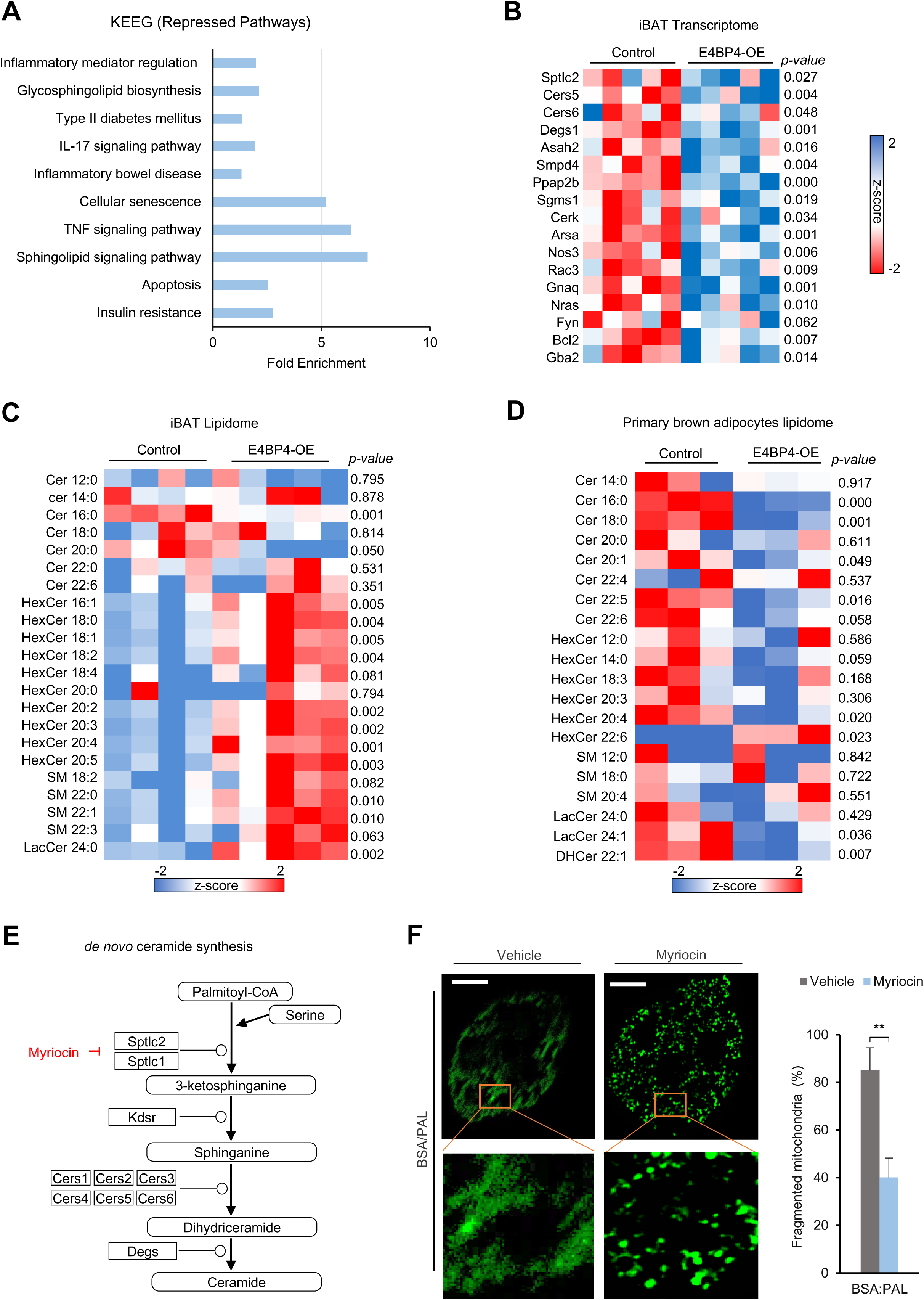
E4BP4 suppresses lipid overload-induced mitochondria fragmentation by repressing ceramide synthesis. A. Gene ontology analysis of repressed KEGG pathways in iBAT from E4BP4-OE versus control mice after 10 weeks of HFD at thermoneutrality (28–29°C). The data are presented as fold-enrichment. B. Heat map of selected genes from the sphingolipid signaling pathway based on RNA-seq analysis of iBAT from E4BP4-OE and control mice (n = 5 per group). Data are represented as Z scores. C. Heat map of iBAT lipidome of control (n=4) and E4BP4-OE (n = 5). Data are represented as Z scores. D. Heatmap showing whole-cell lipidome of Control and E4BP4-OE primary brown adipocytes. Data are represented as Z scores. E. Schematic of de novo ceramide synthesis. F. Left: Images of mitochondrial morphology in primary brown adipocytes by TOM20 immunocytochemistry after 24 h of control treatment (BSA) or palmitate stimulation (BSA:PAL). Representative image. Scale bar, 12 µm. Right: Quantification of cells with a fragmented mitochondrial network (n = 3; 100-180 mitochondria from five images/group/experiment). For F data is presented as mean±SEM. Two-sided P-values were calculated using unpaired Student’s t-tests (B-D–D and F). *P < 0.05, **P < 0.01, ***P < 0.001.

Next, to investigate whether E4BP4 modulates sphingolipid abundance in a cell-autonomous manner, we performed sphingolipidome analysis of primary brown adipocytes pretreated with palmitate (500 µM) for 24h. Our data revealed decreased long-side chain (C16:0, C18:0, C20:1, and C22:5) ceramides, with no changes in HexCer, SM, and LacCer (except HexCer 20:4, HexCer 22:6, and dihydroceramide 22:1) in E4BP4-OE relative to the control (**Figure 4D**). Finally, we investigated whether the inhibition *of de novo* ceramide synthesis could prevent palmitate-induced mitochondrial fragmentation in brown adipocytes. Cells were pretreated with myriocin, an inhibitor of the first and rate-limiting step enzymes serine palmitoyltransferase (Sptlc) isoforms 1 and 2 (**Figure 4E**). Our data showed that myriocin prevented palmitate-induced mitochondrial fragmentation in brown adipocytes (**Figure 4F**). Specifically, myriocin treatment preserved mitochondrial volume and surface area, reduced sphericity, and increased both the number of branches and branch junctions, reflecting maintenance of a more interconnected mitochondrial network (**Figure S4B**). Taken together, these findings suggest that the inhibition *of de novo* ceramide synthesis is sufficient to prevent lipid overload-induced mitochondrial fragmentation in brown adipocytes.

### E4BP4 mitigates Drp1-induced mitochondrial fragmentation by repressing ceramide synthase 6 (Cers6) and *de novo* C16:0 ceramide synthesis in brown adipocytes

Numerous studies have demonstrated that ceramide accumulation leads to reduced mitochondrial oxidative capacity in adipocytes^16,18,19,48^. Of particular interest is C16:0 ceramide, which is derived from the enzymatic activity of Cers5 and Cers6. C16:0 ceramide is a well-known sphingolipid that exerts deleterious effects on mitochondrial oxidative function^49^. Therefore, we investigated whether E4bp4 gain-of-function affects Cers6 expression and C16:0 ceramide levels in iBAT of DIO mice. Accordingly, E4bp4-OE decreased Cer6 mRNA levels (**Figure 5A**) and C16:0 ceramide content compared to those in the control (**Figure 5B**). Next, we employed E4bp4 gain-of-function *in vitro* to analyze *cell-autonomous* repression of C16:0 ceramide synthesis in brown adipocytes. Again, we observed a decrease in Cers6 mRNA levels (**Figure 5C**) and a reduction in C16:0 ceramide content (**Figure 5D**) in palmitate-treated brown adipocytes of the E4BP4 group compared with the control. Ceramides can be formed via *de novo* synthesis or recycling pathways involving sphingomyelin^50^. Hence, to further investigate whether the decreased C16:0 ceramide levels were related to the repression of *de novo* synthesis of ceramides, we pretreated brown adipocytes with a stable palmitic acid (^13^C_16_) isotope (500µM) for 6h and monitored the synthesis of C16:0 ceramides using mass spectrometry (**Figure 5E**). Accordingly, E4bp4-OE reduced *de novo* synthesis of C16:0 ceramide compared to that of the control (**Figure 5E**).

**Figure 5.**
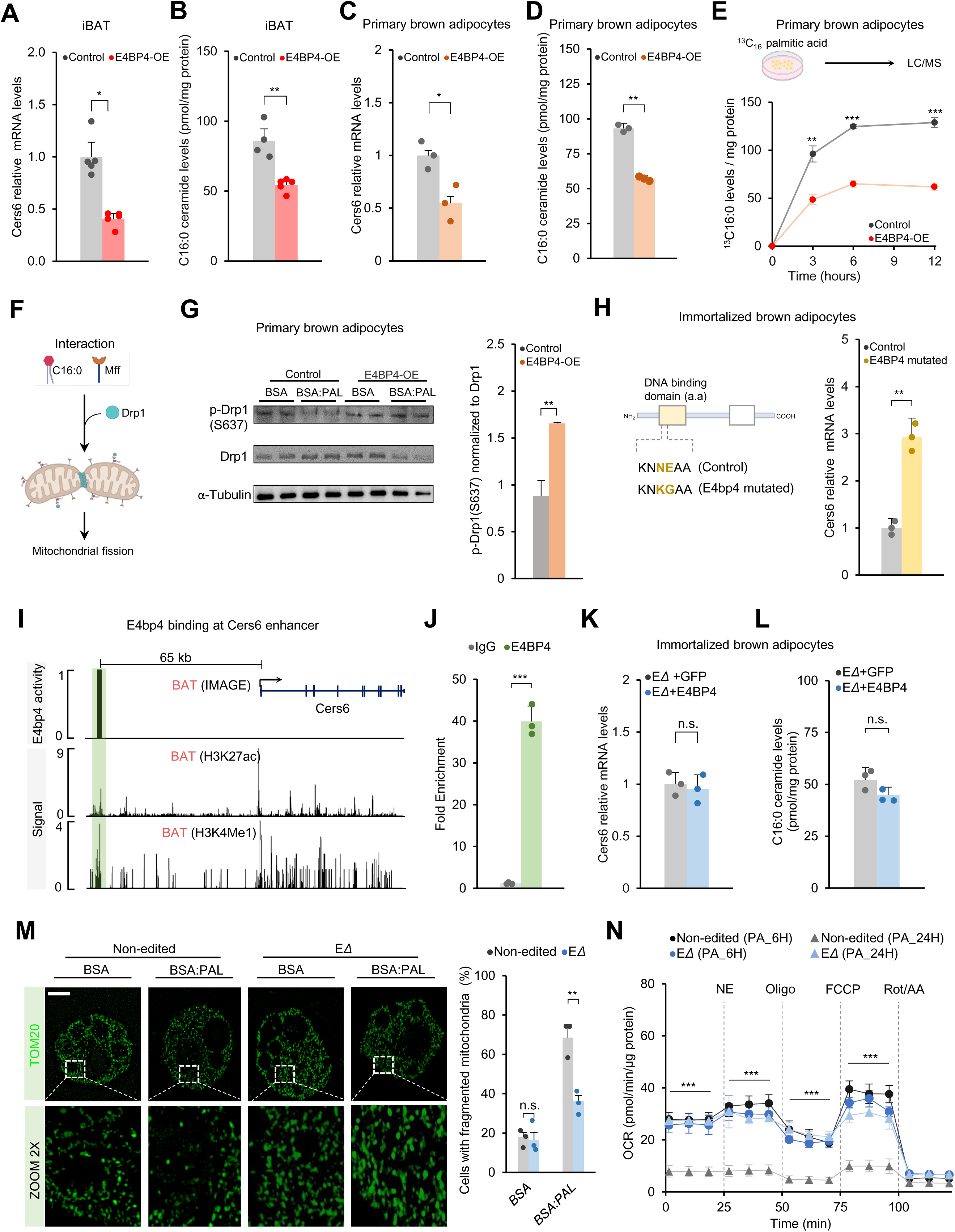
E4BP4 suppresses mitochondrial fragmentation through inhibition of Cers6-driven C16 ceramide synthesis. A. Relative mRNA levels of Cers6 in iBAT of Control and E4BP4-OE mice fed a HFD at 28-29°C. n = 5 per group. B. C16:0 levels in iBAT of Control and E4BP4-OE mice fed a HFD at 28-29°C. n = 5 per group. C. Relative mRNA levels of Cers6 in primary brown adipocytes of the Control and E4BP4-OE groups. n = 3 per group. D. C16:0 quantification by mass spectrometry in primary brown adipocytes of the Control and E4BP4-OE groups. C16:0 levels were normalized to total protein (µg). n = 3 per group. E. C16:0 quantification by mass spectrometry in primary brown adipocytes of Control and E4BP4-OE treated for 3, 6, and 12 h with ^13^C16 palmitic acid. C16:0 levels were normalized to total protein (µg). n = 3 per group. F. A schematic of the mechanism of action of C16:0 binding to the mitochondrial fission factor (Mff), which is a dynamin-related protein-1 (Drp1) adaptor that induces mitochondrial fragmentation. G. Left: Immunoblotting of phospho-Drp1 (S637) and total Drp1 in primary brown adipocytes of Control and E4BP4-OE after 24 h of control treatment (BSA) or palmitate stimulation (BSA:PAL). Representative results are shown from two independent experiments. Gel source data are presented in the Supplementary data. Right: quantification of phospho-Drp1 (S637). H. Left: Schematic of the mutagenesis site directed on the DNA-binding domain or E4bp4. Right: Relative mRNA levels of Cers6 in brown adipocytes from Control and E4BP4-mutated cells. n = 3 per group. I. Top: Peak of E4bp4 activity results from IMAGE analysis in BAT. Bottom: H3K27ac and H3K4me1 signals at *Cers6* loci in BAT. J. Fold enrichment at the binding motif site of E4bp4 (I) in brown adipocytes. The enrichment was assessed by qPCR compared to IgG control. n=3. K. Relative mRNA levels of Cers6 of brown adipocytes mutant E0697815 (ΔE) overexpressing GFP or E4bp4. n=3. L. C16:0 quantification by mass spectrometry of brown adipocytes mutant E0697815 (ΔE) overexpressing GFP or E4bp4. n=3. M. Left: images of mitochondrial morphology in brown adipocytes mutant E0697815 (ΔE) or control (Non-edited) by TOM20 immunocytochemistry after 24 hours of control treatment (BSA) or palmitate stimulation (BSA:PAL). Representative image. Scale bar, 10 µm. Right: quantification of cells with a fragmented mitochondrial network. n = 3. N. The OCR in differentiated brown adipocytes mutant E0697815 (ΔE) or control (Non-edited) after a pre-treatment of 6 or 24 hours with control treatment (BSA) or palmitate stimulation (BSA:PAL). OCR values were normalized by total protein (µg). n = 5 per group. Norepinephrine (NE), oligomycin (oligo), carbonyl cyanide-p-trifluoromethoxyphenylhydrazone (FCCP), and rotenone and antimycin A (Rot/AA). For A-E, G, H, and J-N data are mean ± SEM. Two-sided P values were calculated using unpaired Student’s t-tests (A, B, C, D, E, G, H, J, K, and L). Two-way ANOVA with Tukey’s multiple comparisons test (M and N). *P < 0.05, **P < 0.01, ***P < 0.001.

A previous study demonstrated that C16:0 ceramides interact with Mff in hepatocytes leading to Drp1-induced mitochondrial fragmentation^7^. Thus, we tested whether E4bp4 gain-of-function mitigates mitochondrial fission by modulating Drp1 activity in primary brown adipocytes (**Figure 5F**). Cells were pretreated with palmitate (500 µM) for 24h. Our data revealed that E4bp4-OE enhanced phosphorylation of Drp1^637^, an inhibitory residue, leading to decreased Drp1-mediated mitochondrial fission (**Figure 5G**). Collectively, our data demonstrated that E4bp4 represses Cers6 expression and C16:0 ceramide levels, which in turn prevents mitochondrial fragmentation by reducing Drp1 activity in brown adipocytes.

### E4BP4 is a DNA-binding TF that represses Cers6 by binding to a 65kb upstream enhancer

We sought to determine whether E4bp4 acts as a DNA-binding TF mediating the repression of Cers6 in brown adipocytes. Hence, we used site-directed mutagenesis to alter the amino acid sequence of the E4bp4 bZIP DNA-binding domain, that is, the mutant E4bp4 (**Figure 5H**). Analysis of E4bp4 mRNA levels showed no differences between control versus E4bp4-mutated groups (**Figure S5**). Importantly, our data revealed the loss of Cers6 repression by overexpression of EM E4bp4 in immortalized brown adipocytes, suggesting that E4bp4 acts as a *bona fide* DNA-binding TF (**Figure 5H**).

Next, we explored whether E4bp4 represses Cers6 mRNA expression by directly binding to its regulatory regions. To this end, we analyzed publicly available ATAC-seq and RNA-seq datasets of brown adipocytes using a machine learning-based approach, “*Integrated analysis of motif activity and gene expression changes of transcription factors* (IMAGE)*”* ^51^. Our analysis identified a unique E4bp4 *motif* located 65kb upstream of the Cers6 transcription start site (**Figure 5I**). To evaluate the chromatin status of this region, we analyzed histone H3K27 acetylation (H3K27ac) and H3K4 monomethylation (H3K4me1) in brown fat using the ENCODE data^52^. Accordingly, we observed enriched H3K27ac and H3K4me1 histone marks in this regulatory region, suggesting that this is a typical enhancer site (**Figure 5I**). Moreover, it is worth noting that this enhancer is ENCODE annotated as a distal enhancer “E0697815”. Therefore, for simplicity, we describe this enhancer as “E0697815.” Finally, to validate the binding of E4bp4 to this enhancer, we performed an E4bp4 ChIP-PCR assay. The IgG assay was used as a control. Our data confirmed the enrichment of E4bp4 in E0697815 compared with that in IgG (**Figure 5J**). Altogether, our data demonstrate that E4bp4 acts as a *bona fide* DNA-binding TF to repress Cers6 expression by binding to a 65kb upstream enhancer.

### Deletion of Cers6 upstream enhancer (E0697815) is sufficient to protect mitochondria from palmitate-induced fragmentation

Next, we evaluated the extent to which E0697815 regulates Cers6 expression and C16:0 ceramide levels. To this end, CRISPR/Cas9 was used to alter the genomic sequence of E0697815 in immortalized brown adipocytes (**Figures S3E and S3F**). Sanger sequencing confirmed the presence of an altered nucleotide sequence in the enhancer region (**Figure S3G**). Our data revealed that mutant E0697815 (EΔ) was sufficient to reduce Cers6 mRNA levels (**Figure S3H**) and C16:0 ceramide content (**Figure S3I**) in brown adipocytes, suggesting that E0697815 is a relevant enhancer for controlling Cers6 expression. Moreover, we investigated whether EΔ affects E4BP4-mediated repression of Cers6 expression. Accordingly, we found that the EΔ of Cers6 mitigated E4BP4-induced repression of Cers6 mRNA (**Figure 5K**) and C16:0 ceramide levels (**Figure 5L**).

To further investigate whether the E0697815 mutation in Cers6 was sufficient to prevent palmitate-induced mitochondrial fragmentation, we pretreated the EΔ brown adipocytes with 500µM palmitate for 24h. Mitochondrial morphology was investigated using immunofluorescence assay. As expected, the non-edited control cells displayed fragmented mitochondrial morphology. In contrast, the EΔ of Cers6 was sufficient to prevent palmitate-induced changes in mitochondrial morphology of brown adipocytes (**Figure 5M**). Quantitative analysis demonstrated that EΔ preserved mitochondrial volume and surface area, reduced sphericity, and maintained a higher number of branches and branch junctions, consistent with protection against lipid overload– induced mitochondrial fragmentation (**Figure S4C**). Finally, we conducted OCR analysis to investigate the mitochondrial oxidative capacity in the EΔ Cers6 gene versus non-edited control cells. Brown adipocytes were pretreated with 500µM palmitate for 6 and 24 h before functional analysis. Our data revealed no differences in OCR between EΔ– and control-treated brown adipocytes after 6 h of treatment. However, 24h palmitate treatment significantly decreased OCR at all evaluated respiratory states in the non-edited group, in relation to the EΔ in brown adipocytes. (**Figure 5N**). Taken together, deletion of the 65kb upstream Cers6 enhancer is sufficient to prevent palmitate-induced mitochondrial fragmentation and decrease oxidative capacity in brown adipocytes.

### E4BP4 is co-regulated by PRDM16 to repress Cers6 expression and C16:0 ceramide levels

Given prior evidence that E4BP4 interacts with PRDM16 in beige adipocytes^53^, we sought to determine whether a similar interaction occurs in brown fat. To this end, immortalized brown adipocytes were transfected with Flag-Prdm16 vector, followed by immunoprecipitation. After cell differentiation, Prdm16 was pulled down from the cell extracts using a Flag antibody. Our data revealed the presence of endogenous E4bp4 in the Prdm16 complex (**Figure 6A**). Moreover, we analyzed publicly available Prdm16 ChIP-seq data from iBAT^54^. Interestingly, our analysis revealed a Prdm16 binding site at E0697815 of the Cers6 gene. In contrast, the deletion of Prdm16 (Prdm16KO) abolished the peak at E0697815 (**Figure 6B**).

**Figure 6.**
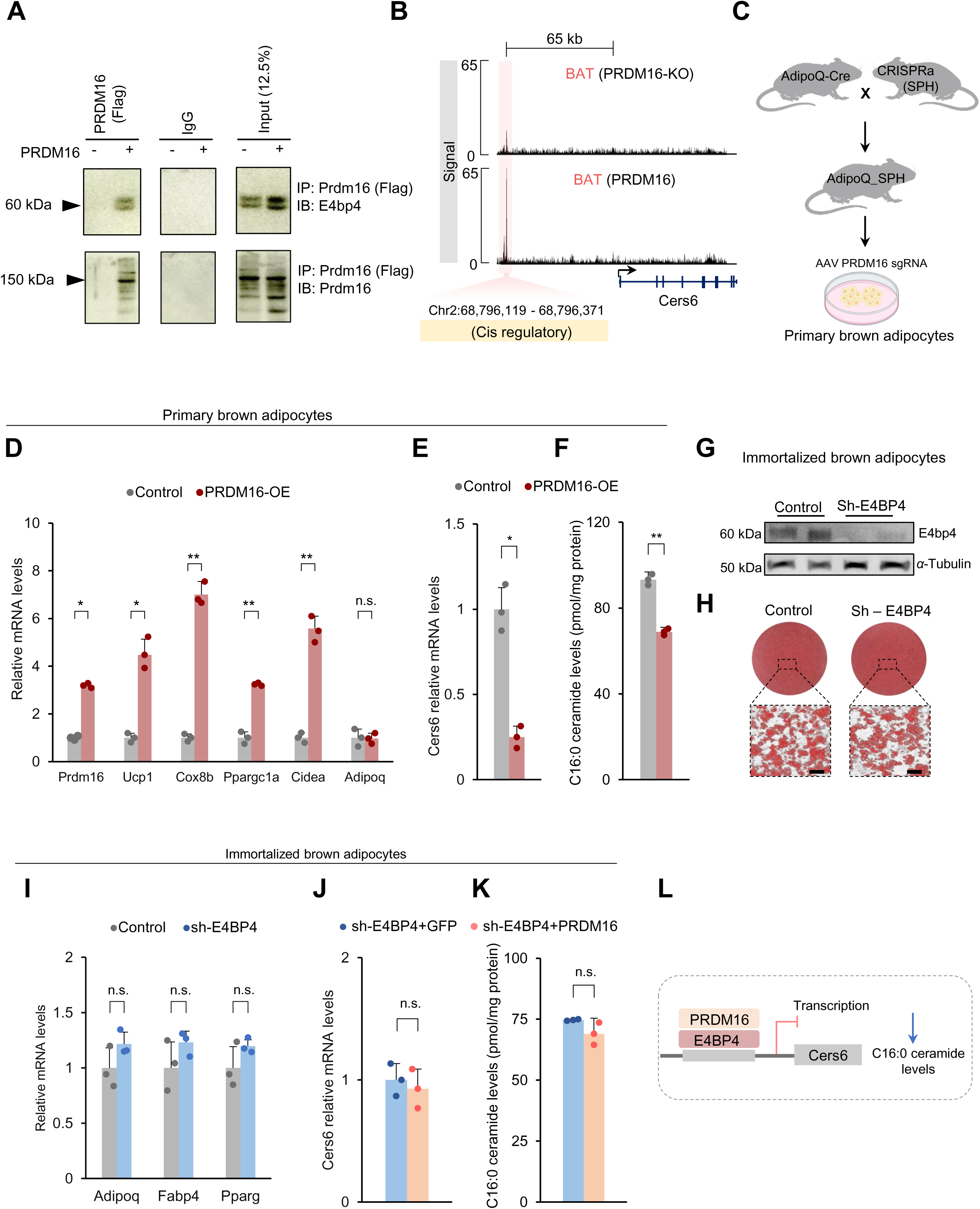
E4BP4 interacts with PRDM16 to inhibit Cers6-driven C16 ceramide synthesis. A. PRDM16 complex was immunopurified from differentiated brown adipocytes. Endogenous E4bp4 was detected by immunoblotting. B. ChIP-seq for PRDM16 and PRDM16 knockout (PRDM16-KO) on the Cers6 gene in BAT. C. Breeding strategy for AdipoQ_SPH mice and littermate control mice. Primary brown adipocytes were used in the experiments. The cartoon was created using a Biorender. D. Relative mRNA levels of indicate genes in primary brown adipocytes of the Control and PRDM16-OE groups. n = 3 per group. E. Relative mRNA levels of Cers6 in primary brown adipocytes of the Control and PRDM16-OE groups. n = 3 per group. F. C16:0 quantification by mass spectrometry in primary brown adipocytes of the Control and PRDM16-OE groups. C16:0 levels were normalized to total protein (µg). n = 3 per group. G. Western blotting for E4bp4 in immortalized brown adipocytes with knockdown for E4bpa (sh-E4BP4) or control. Representative results are shown from two independent experiments. Gel source data are presented in the Supplementary data. H. Representative images of Oil Red O staining in control and sh-E4BP4 brown adipocytes after seven days of differentiation. I. Relative mRNA levels of adipogenic genes in immortalized brown adipocytes of the Control and sh-E4BP4. n = 3 per group. J. Relative mRNA levels of Cers6 of brown adipocytes sh-E4BP4 overexpressing GFP or Prdm16. n=3. K. C16:0 quantification by mass spectrometry of brown adipocytes sh-E4BP4 overexpressing GFP or Prdm16. n=3. L. A proposed mechanism by which E4BP4-PRDM16 complex controls Cers6 expression. For D-F, and I-K data are presented as mean±SEM. Two-sided P-values were calculated using an unpaired Student’s t-test (D, E, F, I, J, and K). *P < 0.05, **P < 0.01, ***P < 0.001.

Previous studies revealed that Prdm16 is involved with the transcriptional repression programs in adipocytes^53,55,56^. Therefore, we investigated whether Prdm16 is associated with the transcriptional repression of Cers6 in brown adipocytes. To this end, we used the CRISPR activation (CRISPRa) mouse model previously established by our group^57^. AdipoQ-Cre mice were crossed with CRISPRa (SPH) mice to establish the conditional expression of CRISPRa machinery in adipose tissues (AdipoQ_SPH). Overexpression of endogenous Prdm16 (Prdm16-OE) was achieved using an AAV single guide RNA (sgRNA) targeting Prdm16 (**Figure 6C**). iBAT was harvested, and SVF was differentiated into brown adipocytes *in vitro*. As expected, Prdm16-OE enhanced the mRNA levels of Prdm16 and thermogenic genes such as Ucp1, Cox8b, Ppargc1α, and Cidea, compared to the control (**Figure 6D**). Moreover, Prdm16-OE led to decreased Cers6 mRNA levels (**Figure 6E**) and C16:0 ceramide content (**Figure 6F**) in palmitate-treated brown adipocytes compared with those in the control. Finally, we investigated whether E4bp4 is necessary for the Prdm16-mediated repression of Cers6 expression. We developed an E4bp4 loss-of-function model by using a lentiviral short hairpin in immortalized brown adipocytes. A scrambled lentiviral sequence was used as the control. Western blot analysis confirmed the loss-of-function of E4bp4 in brown adipocytes (**Figure 6G).** Notably, E4bp4 knockdown did not affect brown adipocyte differentiation, as defined by Oil Red O staining (**Figure 6H**) or expression of adipogenic genes (**Figure 6I**). E4bp4 knockdown mitigated the Prdm16-mediated reduction of Cers6 mRNA levels (**Figure 6J**) and C16:0 ceramide content (**Figure 6K**). Taken together, our data revealed that E4bp4 is co-regulated by Prdm16 to repress Cers6 expression and C16:0 ceramide levels in brown adipocytes (**Figure 6L**).

## DISCUSSION

This study supports a model in which forced E4BP4 expression during DIO prevents mitochondrial fragmentation and oxidative dysfunction in brown fat, at least in part, through repression of ceramide levels in mice. E4BP4 interacts in a transcriptional complex with PRDM16 and directly represses Cers6 mRNA expression and C16:0 ceramide levels by binding to a 65kb upstream Cers6 enhancer. Reduction of Cers6-derived C16:0 ceramide through E4BP4 overexpression or ablation of the Cers6 upstream enhancer was sufficient to prevent palmitate-induced mitochondrial fragmentation and oxidative dysfunction. Notably, the preservation of BAT mitochondrial health by E4BP4 gain-of-function is sufficient to improve systemic glucose homeostasis, independent of weight loss.

E4BP4 is a member of the PAR-bZIP family of transcription factors, characterized by a unique transcriptional repression domain comprising approximately 65 amino acids (residues 299–363) near its C-terminus^32^. Studies using fusion proteins have demonstrated that this specific domain independently acts as a transcriptional repressor when coupled to a heterologous protein, such as the yeast GAL4 DNA-binding domain^30^. These findings prompted us to investigate the transcriptional repression role of E4BP4 in brown fat. In this study, we found that E4BP4 acts as a *bona fide* transcription factor, as evidenced by a mutation in its bZIP domain that disrupts transcriptional repression of Cers6. Recently, the crystal structure of the E4BP4 bZIP domain was elucidated, revealing an α-helical structure that forms dimers via its leucine zipper region and preferentially interacts with the TTACGTAA DNA *motif* ^31^. Notably, disease-associated mutations within the E4BP4’s bZIP domain impair or completely abolish its DNA binding capacity^31^. Additionally, it has been demonstrated that E4BP4 mediates transcriptional repression by recruiting the TBP-binding protein Dr1^58^. However, the mechanisms by which E4BP4 represses Cers6 mRNA remain to be fully elucidated.

We found that E4BP4 is guided by PRDM16 to repress Cers6 mRNA levels by binding to an enhancer region located 65 kb upstream of the gene (E0697815). Notably, the E0697815 mutation abolished the repression of Cers6 mRNA mediated by the E4BP4-PRDM16 complex in brown adipocytes. Furthermore, E4BP4 is necessary for the PRDM16-mediated suppression of both Cers6 mRNA levels and C16:0 ceramide content. These observations reinforce the concept that PRDM16 acts not only as a critical regulator of the thermogenic gene program but also as a broader co-regulator involved in repressing several biological pathways^59^. For example, PRDM16 interacts with GTF2IRD1, a member of the TFII-I family of DNA-binding proteins, to inhibit collagen expression and fibrosis development in adipose tissue^53^. The PRDM16 protein has a short half-life ^60,61^. In this regard, a recent study identified the CUL2-APPBP2 complex as the ubiquitin E3 ligase responsible for controlling PRDM16 stability through polyubiquitination^62^. Inhibition of the CUL2-APPBP2 complex extends the half-life of PRDM16 and mitigates obesity-related metabolic dysfunction^62^. E4BP4 is also tightly regulated at post-translational level^63,64^. However, whether E4BP4 protein half-life can also be extended by inhibition of the CUL2-APPBP2 complex is a matter for future investigation.

Excess nutrients impair mitochondrial oxidative phosphorylation in various cell types, including muscle and pancreatic beta cells^6^. However, it has been hypothesized that under conditions where ATP demand is not the primary driver of oxygen consumption, such as during increased mitochondrial uncoupling in activated brown adipocytes, the negative impact of excess nutrients on mitochondrial oxidative capacity could be minimal^6^. Nevertheless, our findings demonstrated that lipid overload induces mitochondrial fragmentation and oxidative dysfunction in brown fat, at least in part, through the accumulation of C16:0 ceramide. Importantly, the subcellular localization of Cers6 significantly influences mitochondrial morphology and oxidative function, as it is positioned at mitochondria and mitochondria-associated membranes^49^. This localization facilitates the interaction between C16:0 ceramide and Mff, which subsequently activates Drp1-mediated mitochondrial fragmentation^7^. Consistent with these observations, hepatic deletion of Cers6, but not Cers5, improves systemic energy homeostasis in DIO mice^7^. However, it remains possible that reducing C16:0 ceramide levels may enhance the mitochondrial oxidative capacity in brown fat through mechanisms other than regulating mitochondrial morphology. For instance, C16:0 ceramide may affect mitochondrial function by acting on electron transport chain complexes^48,65^. Moreover, recent studies^66,67^ have identified C16:0 ceramide as a ligand for cell membrane G protein-coupled receptors. C16:0 ceramide binds to FPR2, inhibiting thermogenesis via the Gi-cyclic AMP signaling pathway, thereby reducing whole-body energy expenditure^67^. Thus, these are distinct yet non-mutually exclusive mechanisms by which C16:0 ceramide can impair mitochondrial oxidative capacity in adipose tissue.

One of the main findings of our study was that preserving BAT’s mitochondrial morphology and oxidative capacity through E4BP4 gain-of-function improves glucose homeostasis in DIO mice, independent of weight loss. However, we cannot rule out that enhanced energy expenditure plays a role in the improvement of glucose homeostasis, as E4BP4 gain-of-function increases the iBAT temperature and whole-body energy expenditure in DIO mice. The absence of changes in body weight and composition may be explained by a compensatory increase in food intake in the E4BP4 group. Interestingly, these findings align with previous large epidemiological studies, which demonstrated that individuals with detectable BAT had lower circulating glucose levels and approximately a 50% reduced risk of developing type 2 diabetes, irrespective of body weight differences related to their counterparts^68^. Furthermore, interventional studies have shown that activation of BAT using mirabegron, a selective β3AR agonist, significantly improves glucose homeostasis markers in both healthy^69^ and obese insulin-resistant individuals^70^, independent of changes in body weight. These findings reinforce the notion that BAT improves systemic energy homeostasis through mechanisms that extend beyond its classical role of increasing energy expenditure and body weight loss. Therefore, preserving mitochondrial integrity and oxidative capacity should be a primary goal for therapeutic strategies targeting obesity-related metabolic diseases.

### Limitations of the study

We acknowledge the limitations of our lipidomic approach, particularly regarding the analysis of sphingolipids. While our untargeted analysis provided a broad overview of lipid species present in the samples, it may not have fully captured the diversity of sphingolipids, such as sphingomyelins, dihydroceramides, sphingosine, sphinganine, sphingosine-1-phosphate, ceramide-1-phosphate, and gangliosides. To address this, we performed a targeted lipidomic analysis focused on C16:0 ceramide, which was consistently detected across replicates. The downregulation of C16:0 ceramide explains the preservation of mitochondrial morphology (less fragmented) and oxidative capacity observed with the E4BP4 gain-of-function. We also recognize that E4BP4 could improve systemic glucose homeostasis through mechanisms beyond the regulation of ceramide levels. Lastly, we did not examine the effects of E4BP4 in female mice, and the possibility of sexual dimorphism remains to be explored in future studies.

## RESOURCE AVAILABILITY

### Lead contact

Further information and requests for resources and reagents should be directed to and will be fulfilled by lead contact, Carlos Henrique Grossi Sponton.

### Materials availability

Materials generated in this study are available upon request from the lead contact.

### Data and code availability

- Any additional information required to reanalyze the data reported in this work paper is available from the lead contact upon request.
- Uncropped Western Blots are available in Supplementary material.
- This paper does not report original code.
- RNA-seq data have been deposited at GEO and publicly available as of the date of publication.

## Supporting information

reviewers' comments

## ACKNOWLEDGMENTS

We thank Joseane Morari, Marcio Cruz, Erika Roman, and Gerson Ferraz, from the University of Campinas for technical assistance. We thank National Institute of Science and Technology on Photonics Applied to Cell Biology (INFABIC) and Centro Multidisciplinar para Investigação Biológica na Área da Ciência em Animais de Laboratório (CEMIB). We thank the staff of the Life Sciences Core Facility (LaCTAD) from State University of Campinas (UNICAMP) for the genomic analysis. We thank Guilherme Cebinelli from Plataforma de Citometria de fluxo do Hospital Albert Einsten for the Cell sorting analysis. This work used resources of the Centro Nacional de Processamento de Alto Desempenho em São Paulo (CENAPAD-SP). This work was supported by the São Paulo Research Foundation (2020/14725-0, 2020/06057-8, 2021/08354-2, 2013/07607-8, and 2019/15025-5).

## AUTHOR CONTRIBUTIONS

Conceptualization, F. V. R. and C.H.S.; investigation, V.O.F., C.E.L., A.M.Z., M.K.C., F.C.G., A.S., L.R.F., G.L.S., T.G.; writing – original draft, F. V. R. and C.H.S.; writing – review and editing, P.M.M.V., R.F.C., M.A.M., L.A.V. and S.K.; funding acquisition, C.H.S. and L.A.V.; supervision, C.H.S.

## DECLARATION OF GENERATIVE AI AND AI-ASSISTED TECHNOLOGIES

The authors declare no use of generative AI-or AI-assisted technologies in the writing or data interpretation processes of this manuscript. AI-assisted tools were used solely for English proofreading.

## SUPPLEMENTAL INFORMATION

### Figures S1–S5

**Table S1.** FPKM-based quantification of expression changes of transcription factors in interscapular brown adipose tissue (related to Figure 1A).

**Table S2.** Differential gene expression of cold vs thermoneutrality (related to Figure 1E).

**Table S3.** Differential gene expression of Control vs E4BP4-OE (related to Figure 4).

**Table S4.** Primer sequences and probes for qRT-PCR.

**Video S1.** Live-cell fluorescence of MitoDendra2 in primary brown adipocytes (related to Figure 3G).

## FIGURE TITLES AND LEGENDS

**Figure S1.**
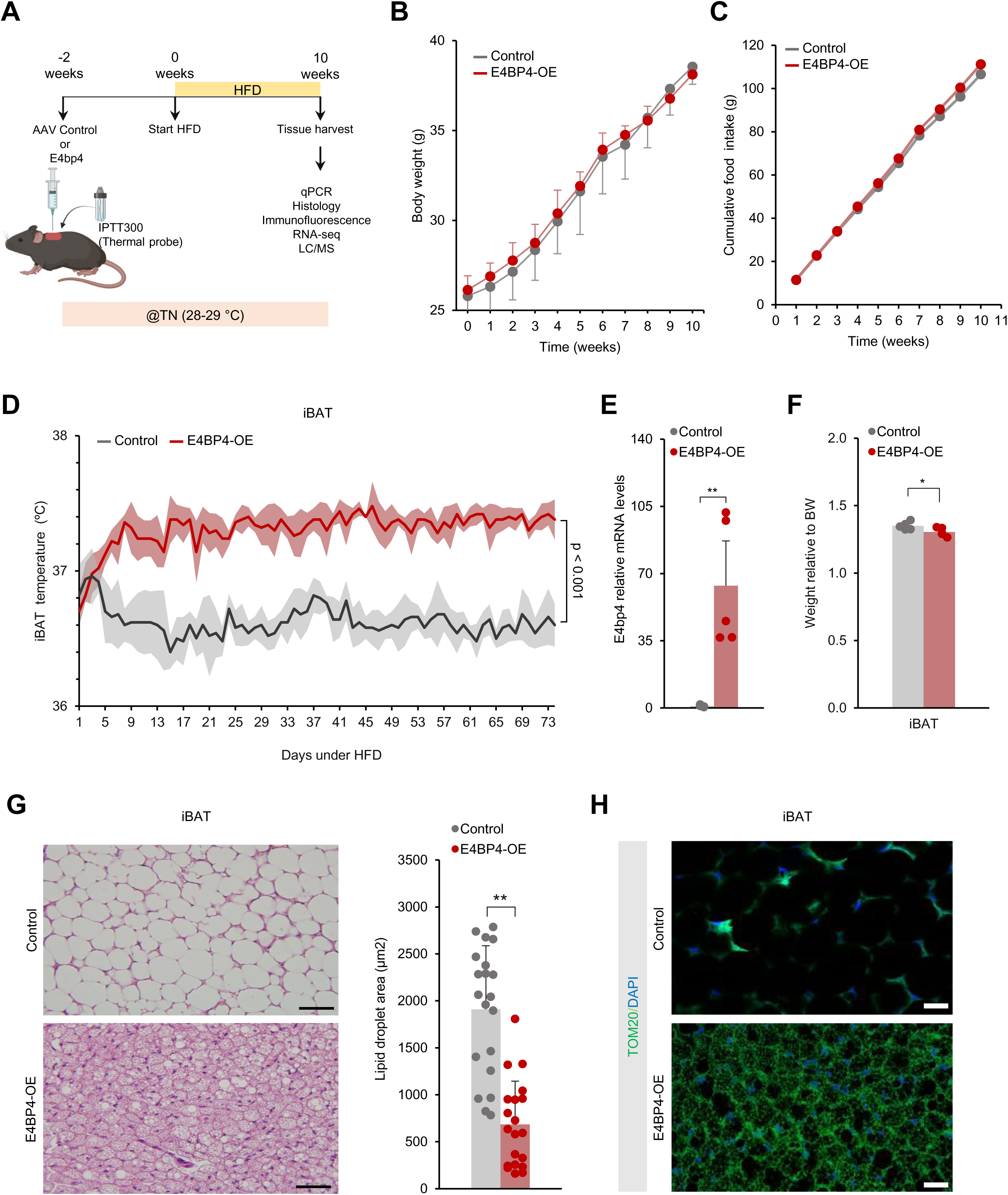
Analysis of E4BP4-OE in DIO mice using AAV constitutive CMV promoter (Related to figure 2) A. Schematic illustration of experimental design (A-H) for overexpression of E4bp4 (E4BP4-OE) or control in iBAT. The cartoon was created using a Biorender. B. Body-weight of Control or E4BP4-OE mice under HFD at 28-29 °C. n = 5 per group. C. Cummulative food intake of Control or E4BP4-OE mice under HFD at 28-29 °C. n = 5 per group. D. Temperature of iBAT of Control or E4BP4-OE mice under HFD at 28-29 °C. n = 5 per group. E. Relative mRNA levels of E4pb4 in iBAT of Control and E4BP4-OE mice fed a HFD at 28-29°C. n = 5 per group. F. Tissue weight normalized to body weight of Control and E4BP4-OE mice fed a HFD at 28-29°C. n = 5 per group. G. Left: representative images of Hematoxylin and eosin (H&E) staining of iBAT from control and E4BP4-OE mice fed a HFD at 28-29°C. Scale bar: 50μm. Right: Lipid droplet area quantification. n = 3 per group. H. Images of mitochondrial iBAT of Control and E4BP4-OE by TOM20 immunocytochemistry Representative image. Scale bar, 50 µm. For B-G data are presented as mean±SEM. Two-sided P-values were calculated using an unpaired Student’s t-test (B, C, E, F, and G). Two-way ANOVA with Tukey’s multiple comparisons test (D). *P < 0.05, **P < 0.01, ***P < 0.001.

**Figure S2.**
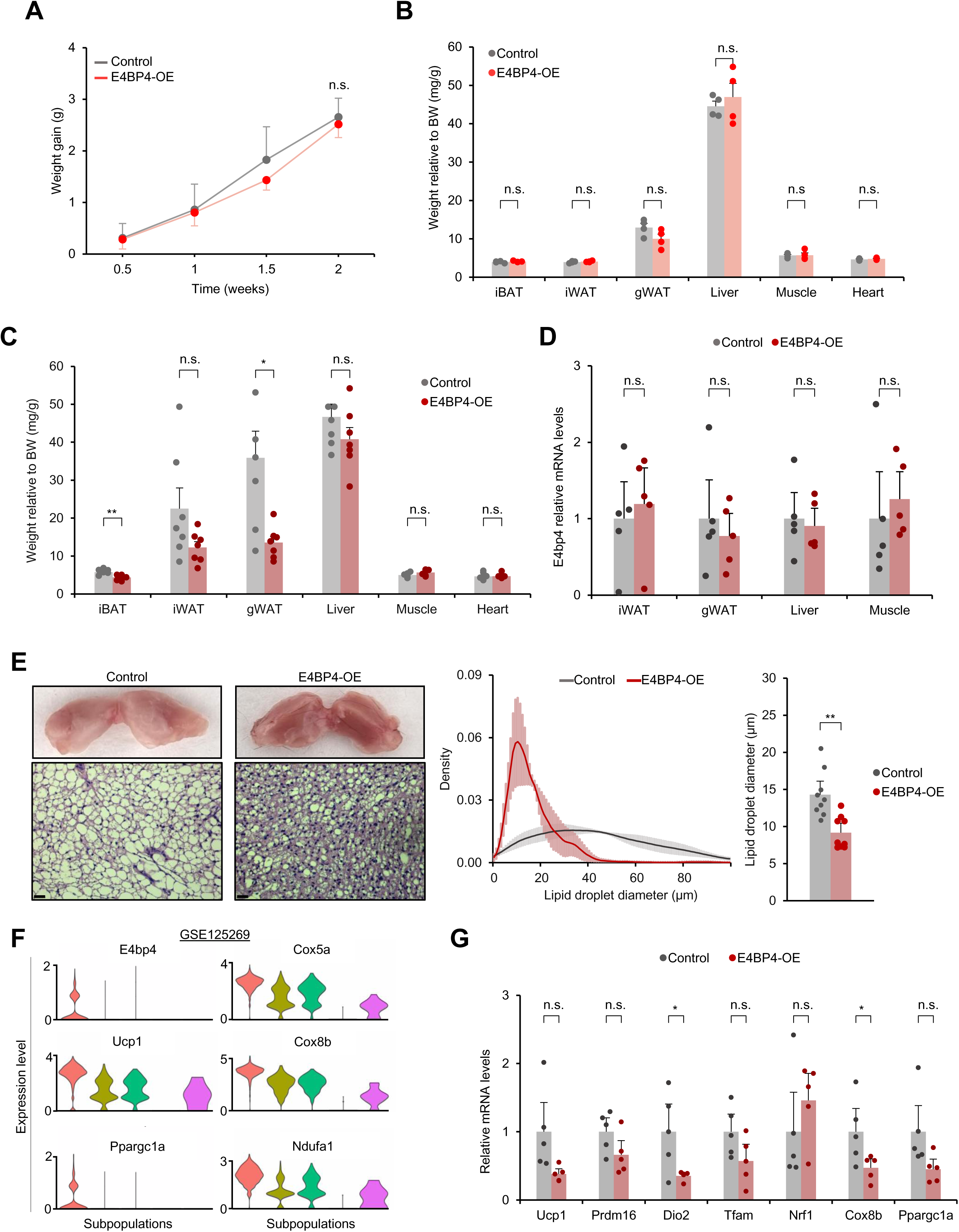
Analysis of E4BP4-OE in DIO mice using AAV adiponectin promoter (Related to Figure 2) A. Weight of E4BP4-OE or control mice under HFD at 28-29 °C during 2 weeks. n = 4 per group. B. Tissue weight normalized to body weight of Control and E4BP4-OE mice fed a HFD at 28-29°C during 2 weeks. n = 4 per group. C. Tissue weight normalized to body weight of Control and E4BP4-OE mice fed a HFD at 28-29°C during 10 weeks. n = 7 per group. D. Relative mRNA levels of E4pb4 in the indicate tissues of Control and E4BP4-OE mice fed a HFD at 28-29°C during 10 weeks. n = 7 per group. E. Left: representative images of Hematoxylin and eosin (H&E) staining of iBAT from control and E4BP4-OE mice fed a HFD at 28-29°C during 10 weeks. Scale bar: 50μm. Right: lipid droplet diameter distribution and quantification. n = 3 per group. F. Violin plots showing the distribution of normalized expression values of indicated genes across cells of the adipocyte clusters. G. Relative mRNA levels of indicate genes of Control and E4BP4-OE mice fed a HFD at 28-29°C during 10 weeks. n = 7 per group. For A-E, and G data are presented as mean±SEM. Two-sided P-values were calculated using an unpaired Student’s t-test (A-E, and G). *P < 0.05, **P < 0.01, ***P < 0.001.

**Figure S3.**
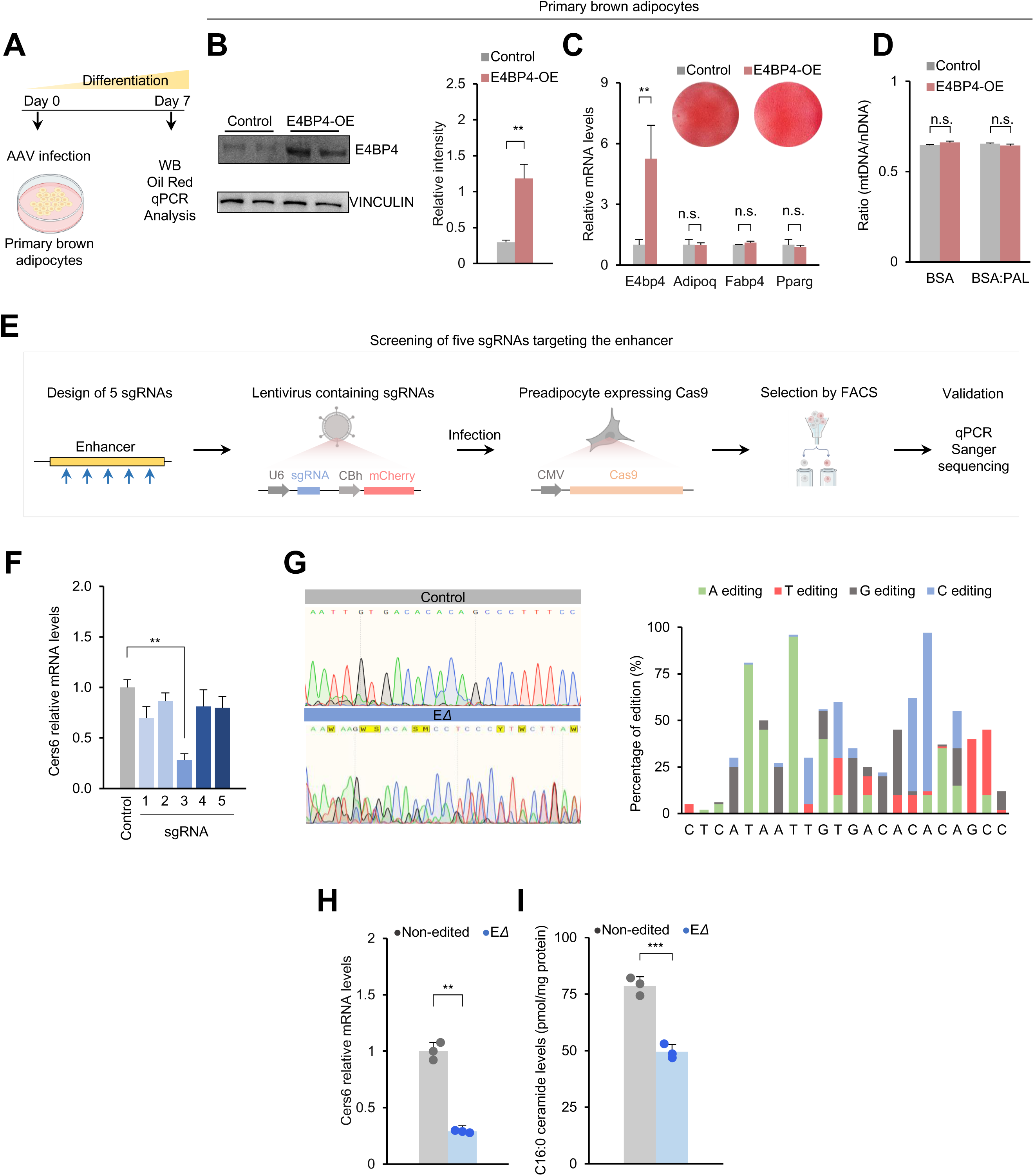
E4bp4 gain-of-function and mutant (ΔE) Cers6 enhancer (E0697815) models in brown adipocytes (Related to Figure 3 and 5) A. Schematic illustration of experimental design (B-D) for overexpression of E4bp4 (E4BP4-OE) or control in primary brown adipocytes. The cartoon was created using a Biorender. B. Left: western blotting for E4bp4 in in primary brown adipocytes of Control and E4BP4-OE. Representative results are shown from two independent experiments. Gel source data are presented in the Supplementary data. Right: quantification of western blot. C. Relative mRNA levels of indicate genes in primary brown adipocytes of the Control and E4BP4-OE groups. n = 3 per group. D. Ratio of mitochondrial DNA (mtDNA) and nuclear DNA (nDNA) in primary brown adipocytes of the Control and E4BP4-OE groups. n = 3 per group. E. Schematic illustration of experimental design for the screening of sgRNAs targeting the enhancer (F-G). The cartoon was created using a Biorender. F. Relative mRNA levels of Cers6 in brown adipocytes mutant E0697815 (ΔE) for the Validation of sgRNA. n = 3 per group. G. Left: Sequencing analysis of CRISPR/Cas9-mediated enhancer E0697815 (ΔE) ablation in brown adipocytes. Nucleotide sequences of targeted enhancer of cells treated with sgRNA and control. Right: percentage of nucleotide mutations derived from Sanger sequencing data (G). H. Relative mRNA levels of Cers6 in brown adipocytes mutant E0697815 (ΔE) or Control. n=3 per group. I. C16:0 quantification by mass spectrometry in brown adipocytes mutant E0697815 (ΔE) or Control. n=3 per group. For B, C, D, F, H, and I data are presented as mean±SEM. Two-sided P-values were calculated using an unpaired Student’s t-test (B, C, D, F, H, and I). *P < 0.05, **P < 0.01, ***P < 0.001.

**Figure S4.**
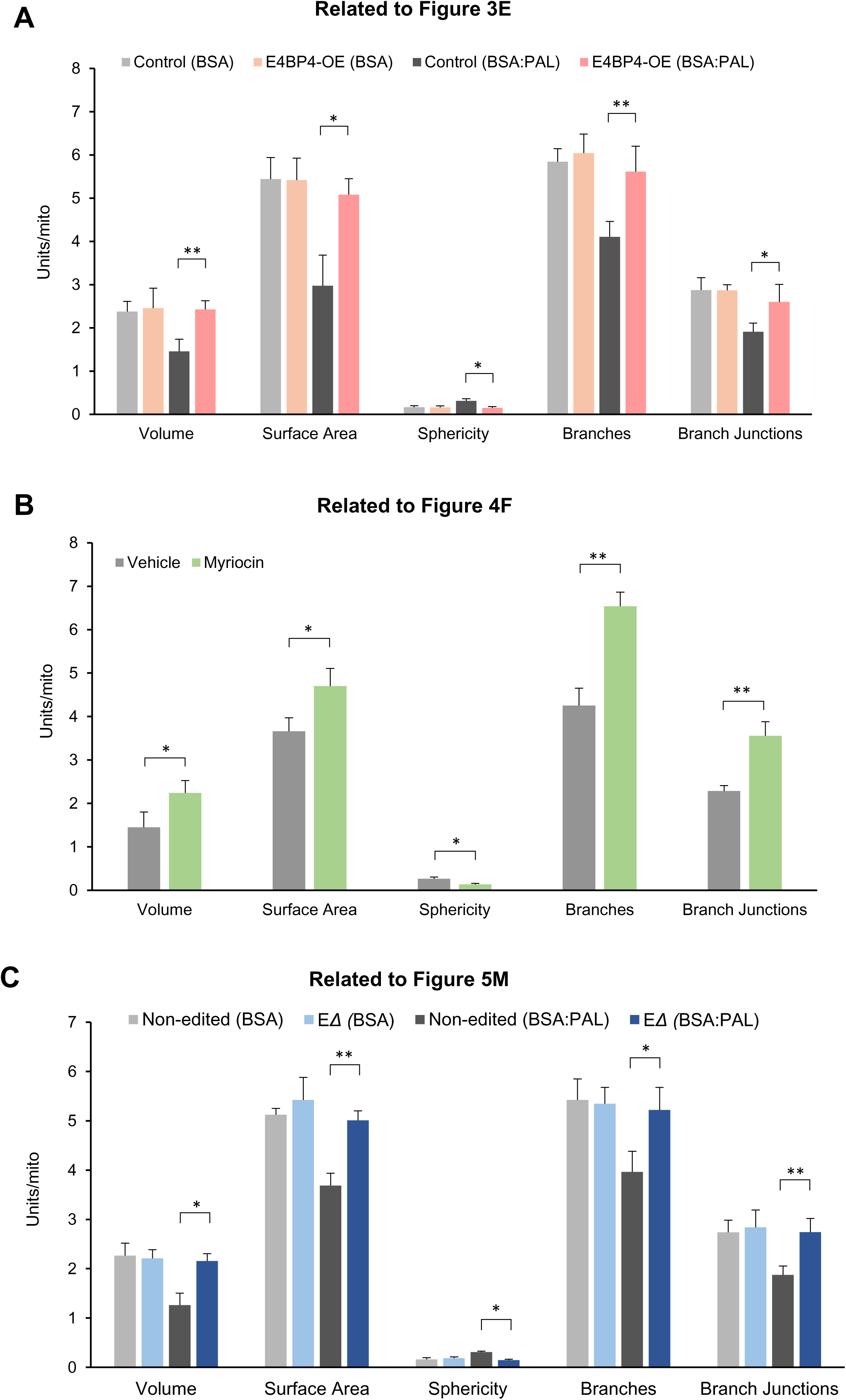
Mitochondrial morphology characterization. A. Quantification of mitochondrial morphology. Mean mitochondrial volume (units in μm3), Mean mitochondrial surface area (units in μm^2^), mitochondrial sphericity index, mean number of branches and junctions per mitochondrion. B. Quantification of mitochondrial morphology. Mean mitochondrial volume (units in μm3), Mean mitochondrial surface area (units in μm^2^), mitochondrial sphericity index, mean number of branches and junctions per mitochondrion. C. Quantification of mitochondrial morphology. Mean mitochondrial volume (units in μm3), Mean mitochondrial surface area (units in μm^2^), mitochondrial sphericity index, mean number of branches and junctions per mitochondrion. For A, B, and C data are presented as mean±SEM. Two-sided P-values were calculated using an unpaired Student’s t-test. *P < 0.05, **P < 0.01, ***P < 0.001.

**Figure S5.**
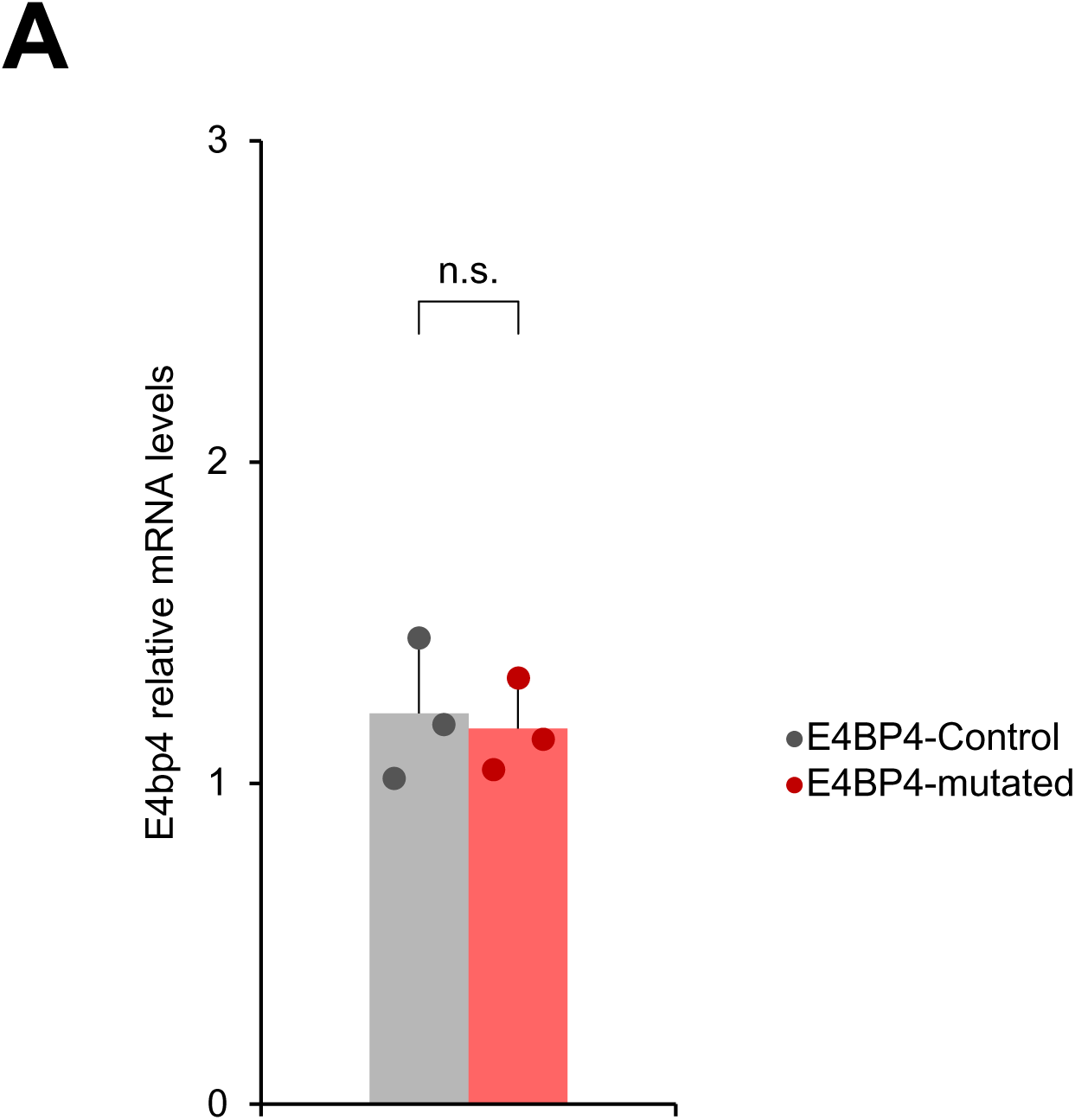
Quantification of E4bp4 mRNA levels. **A.** Relative mRNA levels of E4bp4 in brown adipocytes from E4BP4 and E4BP4-mutated cells. n = 3 per group. data are presented as mean±SEM. Two-sided P-values were calculated using an unpaired Student’s t-test.

## EXPERIMENTAL MODEL AND STUDY PARTICIPANT DETAILS

### Mouse husbandry

All animal experiments were performed in compliance with protocols approved by the University of Campinas Guide for the Care and Use of Laboratory Animals (protocol CEUA # 6102-1/2022; #6619-1/2025). All mice were housed under a 12 h light – 12 h dark cycle. Room temperature mice were housed at 22 °C. Mice kept under thermoneutral conditions were placed inside an incubator maintained at a temperature of 28-29 °C. Mice were fed a standard diet or high-fat diet (HFD; 60% kcal from fat) and had free access to food and water. Mice were fasted for 6 h prior to metabolic measurements and brown adipose tissue respiration. SPH mice were acquired from Jackson Laboratory (Strain# 031645) and crossed with Adipo-Cre (Jackson Laboratory, Strain# 028020). To generate reporter primary brown adipocytes for mitochondria, B6;129S-Gt(ROSA)2^6Sortm1(CAG-COX8A/Dendra2)Dcc^/J (JAX stock #018385) mice were crossed with mice expressing Cre recombinase under the control of the adiponectin (Adipoq) promoter. This breeding strategy yielded mice expressing Dendra2, a photoactivatable protein located in the mitochondria (PhAM) of adipocytes, as well as Cre-negative littermate controls. Genotyping was performed using the specific primers listed in Table S4, and representative PCR gel images are provided in the Supplementary Materials. Male mice were used unless otherwise stated.

### Primary cell isolation and culture

Brown adipose tissue (BAT) was dissected and digested with dispase II (Sigma D4693) and collagenase D (Sigma 11088882001) at 37 °C for 25 min with agitation. The stromal vascular fraction (SVF) containing preadipocytes was separated from the mature adipocytes by centrifugation at 700 × g for 10 min. SVF cells were then plated on coated 6 well plates in culture medium containing Dulbecco’s modified Eagle’s medium (DMEM) (Thermo Fisher, 10569044) supplemented with 10% fetal bovine serum (FBS) and 1% penicillin/streptomycin (P/S) (Gibco, 15140). Immortalized brown preadipocytes kindly donated by Dr. Marcelo Mori (University of Campinas) were maintained in (DMEM) supplemented with 4.5 g/L glucose, 10% FBS, and 1% P/S at 37 °C with 5% CO2. Brown adipocyte differentiation was induced by treating the confluent preadipocytes with DMEM containing 10% FBS, 0.5 mM isobutylmethylxanthine, 125 μM indomethacin, 2 μg/ml dexamethasone, 20 nM insulin, 1 nM T3, and 1 μg/ml rosiglitazone. Two days after induction, the cells were switched to maintenance medium containing 10% FBS, 20 nM insulin, 1 nM T3, and 1 μg/ml rosiglitazone. For Adeno-X 293 cells, Lenti-X 293T cells were grown and maintained in DMEM containing 10% FBS and 1% P/S.

### Experimental Details

#### Plasmid construction and acquisition

To overexpress E4BP4 under a constitutive promoter, the plasmid carrying the GFP protein (Addgene catalog no. 49055) was replaced with the sequence of E4BP4 amplified from the plasmid (Addgene catalog no. 34572) using Platinum SuperFi II DNA Polymerase (Thermo Fisher). After ligation, specific primers from the promoter and E4BP4 sequence were used to confirm insertion. To overexpress E4BP4 under the control of the adiponectin promoter, the sequence of E4BP4 amplified from the plasmid (Addgene catalog no. 34572) using Platinum SuperFi II DNA Polymerase (ThermoFisher) was inserted into the MCS region of pAAV-ADP-MCS-FLAG (Addgene catalog no. 192360). After ligation, specific primers from the promoter and E4BP4 sequence were used to confirm insertion. PRDM16 overexpression was achieved using a retroviral plasmid purchased from Addgene (catalog no. 15504).

### Lentivirus packaging

For viral production, Lenti-X 293T packaging cells were transfected with DNA constructs using pCMV delta R8.2 (Addgene, catalog no.12263) and pMD2.G (Addgene, catalog no. 12259) using polyethylenimine (Sigma-Aldrich, 408727). After 48 h, the viral supernatant was collected and filtered through a 0.45 mm filter.

### Adeno-Associated Virus (AAV) packaging and titration

AAV packaging and titration were performed, as previously described^57^. In brief, AAV was produced by triple transfection of a targeting vector plasmid with pAdDeltaF6 plasmid (Addgene catalog no. 112867) and pAAV2/8 plasmid (Addgene catalog no. 112864) into Adeno-X 293 cells (Takarabio) using polyethylenimine (1 µg/µl) (Sigma-Aldrich, 408727). The cells were collected 72 h post-transfection and purified using an Amicon Ultra-0.5 centrifugal filter (Merck Millipore, UFC510024).

### Human brown adipocytes

Human brown preadipocytes were maintained in Dulbecco’s modified Eagle’s medium F12 (DMEM-F12) supplemented with 4.5 g/L glucose, 10% fetal bovine serum (FBS), and 1% penicillin/streptomycin at 37 °C with 5% CO2. Brown adipocyte differentiation was performed as previously described^71^. In brief, cells were pretreated with 3.3 nM BMP7 for 6 days prior to induction. Immortalized human preadipocytes were cultured in high-glucose DMEM supplemented with 10% FBS and antibiotics at 37°C with 5% CO₂. Upon confluency, differentiation was induced using a defined medium containing insulin, T₃, dexamethasone, IBMX, indomethacin, biotin, and pantothenate. Medium was refreshed every 3 days for 12 days. After differentiation, the cells were treated with forskolin (20 µM) or vehicle for 24h.

### Gain-of-function in immortalized brown adipocytes

E4bp4 (Addgene, catalog no. 34572) and Prdm16 (Addgene, catalog no. 15503) plasmids were transfected using Lipofectamine 3000 (Thermo Fisher Scientific) according to the manufacturer’s protocol. Briefly, preadipocytes at 80% confluence were transfected, and 24 h after transfection, the cells were switched to differentiation medium.

### AAV overexpression in primary brown adipocytes

Primary brown adipocytes were infected with AAV expressing E4BP4 (1×10^10^ vg/ml) in the presence of polybrene (8 ng/μl) (Sigma-Aldrich, H9268), and 18 h after transduction, the cells were switched to differentiation medium. The analyses were performed on mature adipocytes.

### Loss-of-function of E4bp4

The short hairpin (shRNA) sequence was cloned into an Age1– and EcorI-digested PlKO.1 hygro plasmid (Addgene, catalog no. 24150) by annealing oligos sh-E4bp4 described in Table S4. Generation of the sh-E4bp4 cell line using lentiviral transduction. 9B cells were transduced with viral supernatant in the presence of polybrene (8 ng/μl) (Sigma-Aldrich, H9268). Antibiotic selection with hygromycin was initiated 5 days after transduction.

### Genome editing of enhancer of cers6

Five sgRNAs target sites were predicted using the CRISPOR tool available in Genome (https://genome.ucsc.edu). Oligos (Table S4) were annealed and cloned into a BsmBI-digested sg-shuttle-RFP657 vector (Addgene, catalog no. 134968). WT1-KO cells were transduced with viral supernatant in the presence of polybrene (8 ng/μl) (Sigma-Aldrich, H9268). After 5 days of expression, positive cells for RFP were isolated by fluorescence (excitation/emission at 611/657 nm) cell sorting, and individual clones were expanded and tested for loss of function of the enhancer of Cers6 using qPCR. To validate the mutation, we first amplified the enhancer by PCR, and the gel band was subjected to Sanger sequencing. To confirm the mutation, sequencing peaks were analyzed using Edit R^72^.

### Palmitate treatment

Palmitate treatment was performed as previously described^7^. In brief, 500 μM palmitic acid (Sigma-Aldrich, P0500) was conjugated to 1% (w/v) fatty-acid-free low-endotoxin BSA (Sigma-Aldrich, A8806) in DMEM at 37°C for 25 min. Following this, the cells were incubated with BSA-conjugated PAL or BSA for 6 or 24 h at 37°C with 5% CO2.

### Measurement of c16 flux using stable isotopes

Fully differentiated primary brown adipocytes in a 6-well plate were switched to DMEM containing 1% FBS and 500 µM ^13^C16 palmitate:BSA (Sigma-Aldrich, 605573) for 3, 6, and 12 h. Lipids were extracted and analyzed using mass spectrometry. Protein concentration was determined using the Pierce BCA Protein Assay Kit (Thermo Fisher Scientific, 23225) and used to normalize the metabolites.

### Measurement of oxygen consumption rate

The oxygen consumption rate (OCR) was measured with the Seahorse XFe Extracellular Flux Analyzer (Agilent) in a 24-well plate, as previously described^19^, with some modifications. Briefly, primary brown adipocytes were transduced with AAV-E4BP4 or AAV-GFP, and 18 h later, the cells were switched to differentiation medium. The cells were trypsinized on day 8 and plated into 24-well seahorse plates as previously described. The experiment was performed in seahorse assay media supplemented with 10 mM glucose, 2 mM glutamine, and 1 mM pyruvate. Subsequently, drug injections were administered in the following sequence: noradrenaline (1 µM), oligomycin (5 µM), FCCP (5 µM), and Rotenone + Antimycin (1/2 µM). The same methodology was applied to immortalized brown adipocytes with enhancer mutations.

### Immunohistochemistry

Cover slips with fully differentiated adipocytes were fixed using 4% PFA in 0.1 M PBS for 15 min at room temperature. After two PBS washes, the cells were incubated in blocking solution (10% normal donkey serum in 0.1 M PBS containing 0.2% Triton X-100) for 1.5 hours at room temperature. Next, the adipocytes were incubated overnight at 4°C in fresh blocking solution (3% normal donkey serum) containing rabbit anti-TOM20 (1:200; MA5-24859, Thermo Fisher Scientific) without mixing. After three PBS washes, the cells were incubated in a blocking solution containing donkey anti-rabbit FITC (1:500; ab6798 or Ab6941) for 90 min at room temperature. The slides were washed three more times with PBS, mounted, and imaged using a Zeiss LSM880 Airyscan microscope with the ZEN Software (Carl Zeiss). Immunopositive cell counts were averaged from three cover slips per experimental replicate, with 4-5 fields imaged per cover slip. Quantification of immunopositive cells per image was performed using Fiji software, employing the Deconvolution plugin (Deconvolved 3D) for mitochondrial isolation and the Mitochondria Analyzer plugin for quantification. Cells with significant mitochondrial network fragmentation were counted as described previously^7^, and image representation was generated from the maximum projection of z-stacks.

### High-resolution time-lapse imaging and mitochondrial signal quantification

Live-cell super-resolution images were acquired using a Zeiss LSM880 confocal microscope equipped with an Airyscan detector (Carl Zeiss AG, Germany) using a Plan-Apochromat 63×/1.4 oil immersion magnification. All time-lapse acquisitions were conducted under physiological conditions using a stage-top incubation system that maintained cells at 37 °C in a humidified atmosphere with 5% CO₂. Dendra2 was excited in its native state using a 488 nm laser line and in its photoconverted state using a 543 nm laser. Photoconversion was induced by illuminating a defined region of interest with a 405 nm laser, 5%, 90 iterations for bleaching, with a scan speed of 6. The time series consisted of 9 cycles with a frame interval of 30 min. The 3D images were 1024 × 1024, 8 bits, zoom 3, resolution 4.8 px/μm. Image analysis was performed using the FIJI/ImageJ software. Single-channel images were processed by generating maximum-intensity Z-projections, when applicable. The background was subtracted using a rolling ball radius of 30 pixels followed by a Gaussian filter. Pixel intensity values were restricted to a range of 100–350 to exclude background noise and automatic thresholding (default, dark background) was applied. Binary masks were generated and used to quantify area fraction. Logical operations (AND) between processed images were employed to isolate the signal overlap between green and red images; thus, the fusion of photoconverted mitochondria with any mitochondria from the adipocyte network. The fusion process is then measured using the resulting masks. Co-localization was normalized using the ratio of the number of interacting particles to the total number of particles per area.

### Time-lapse microscopy analysis

For the time-lapse analysis, we used the methodology described previously ^45^, with some modifications. In brief, differentiated adipocytes were pretreated for 24 h with BSA, palmitate, or BSA before treatment with fluorescent dyes. Mitotracker Green (MTG) was used at 200 nM for 90 minutes and subsequently removed by washing before imaging. Tetramethylrhodamine-ethyl-ester-perchlorate (TMRE) was loaded at 30 nM and remained present during the imaging. The initial photograph was captured at this time point (−1). Subsequently, cells were treated with PAL, BSA, or BSA, initiating a time-lapse sequence for 60 min. Time-lapse images were captured using a Zeiss LSM880 Airyscan microscope with the ZEN Software (Carl Zeiss). The adipocytes were maintained at 37°C in 5% CO2.

### Confocal microscopy image processing and analysis

Mitochondrial images were analyzed using Fiji software. The images were smoothed and binarized to analyze the area of single particles. The binary images were then skeletonized, and each skeleton was quantified to obtain the number of networks, junctions, and branches, as well as the volume, surface area, sphericity of each particle. Cells exhibiting significant fragmentation of the mitochondrial network were quantified, as previously described^7^.

### Oil red o staining

After the cells were fixed in 4% PBS-buffered formaldehyde for 30 min, they were washed 3 times and stained with ORO solution (0.3% ORO in 60% isopropanol) for 20 min.

### Protein analysis

Cell lysates and tissues were lysed in RIPA buffer (50 mM Tris (Merck), pH 8, 150 mM NaCl (Merck), 5 mM EDTA (Merck), 0.1% w/v SDS (Carl Roth), 1% w/v IGEPALCA-630 (Sigma-Aldrich), 0.5% w/v sodium deoxycholate (Sigma-Aldrich)) freshly supplemented with protease inhibitors (Sigma-Aldrich) and phosphatase inhibitors (Roche). The cell lysates were centrifuged for 20 min (4 °C and 13,000 × *g*). Protein concentrations were determined using the Pierce BCA assay (Thermo Fisher Scientific), following the manufacturer’s instructions. Equal amounts of solubilized proteins were loaded onto SDS-PAGE gels and blotted onto nitrocellulose or PVDF membranes (Bio-Rad). The membranes were incubated overnight with the primary antibody (E4BP4, 1:1000) in 5% BSA-TBST at 4 °C. After washing with TBST ((200 mM Tris (Merck), 1.36 mM NaCl (Merck), 0.1% v/v Tween20 (Sigma-Aldrich)), the membranes were incubated in secondary antibody (Santa Cruz) solutions (1:10,000 in 3% BSA) for 1-2 h at RT and revealed using an enhanced chemiluminescence reagent (Clarity Max Western ECL Substrate, BioRad). Full scans of the uncropped blots are available in Data S2. and incubated overnight with the primary antibodies, as detailed in the reagent table.

### AAV injection

Mice were anesthetized with 1-2% isoflurane using a Vaporizer AI-100 (Insightltda). A 0.3-0.8 cm longitudinal incision was made in the skin at the interscapular region to expose the brown adipose tissue. Thirty microliters of AAV (2×10^11^ vg/ml) was administered to both lobes of the brown fat depot of mice aged 6 weeks using a Hamilton syringe. Following the injections, IPTT-300 Temperature Transponders (PLEXX, 11059) were inserted at the midline of the interscapular region. All surgical procedures were performed at least two weeks prior to the initiation of the experiment to allow full recovery.

### Animals under cold exposure

Wild-type mice (C57BL6J) were maintained on a 12-h light–dark cycle. For cold exposure, mice were housed at 4°C for 0–14 days. All in vivo experimental procedures in this study were performed during the light phase (07:00–10:00 AM) to avoid significant data variability due to circadian rhythm fluctuations^73^.

### Indirect calorimetry

Metabolic measurements were obtained using an LE405 Gas Analyzer (Panlab, Harvard Apparatus, Holliston, MA, USA) indirect calorimetry system. Food and water were provided ad libitum to the appropriate devices and were measured using built-in automated instruments. The animals were allowed to acclimatize to the cages for at least 18 h before data acquisition. Data analysis was performed using CalR version 2.0^74^.

### Brown adipose tissue histology

Dissected BAT samples were fixed in 4% paraformaldehyde, embedded in paraffin, sectioned at 5 mm, and stained with hematoxylin and eosin (H&E). To visualize lipid deposition H&E-stained sections were imaged on a Zeiss Axio Observer microscope (Carl Zeiss). The diameter and area were calculated using ImageJ software. The diameter distribution was estimated for each tissue using weighted kernel density estimation using the Seaborn library^75^.

### Brown adipose tissue immunohistochemistry

Dissected BAT samples were fixed with 4% paraformaldehyde, embedded in paraffin, and sectioned. First, we employed a method of deparaffinization with 100% xylene for 5 min at room temperature, and a second incubation with fresh 100% xylene for an additional 5 min to ensure complete removal of paraffin. Next, the slides were rehydrated with 100% ethanol for 5 min and sequentially transferred through a series of ethanol solutions with decreasing concentrations (95%, 90%, 80%, and 70%), and immersed in each solution for 5 min to gradually rehydrate the tissue sections. Finally, the slides were resuspended in PBS. For staining, the same methods as for immunohistochemistry using anti-Tom20 (used for cells) were used without modifications. Images were obtained using a Zeiss LSM880 Airyscan microscope with the ZEN Software (Carl Zeiss).

### Brown adipose tissue respiration

Tissue respiration was performed using an O2K high-resolution respirometer (Oroboros Oxygraph-2k; Innsbruck, Austria). Freshly isolated tissues were weighed 2-3 mg and homogenized with BIOPS buffer (2.77 mM of CaK2EGTA, 7.23 mM K2EGTA, 0.1 µM free calcium, 20 mM imidazole, 50 mM K+/MES, 0.5 mM dithiothreitol, 7 mM MgCl2) and phosphocreatine (15 mM, pH 7.1) at 0-4°C. The homogenized tissue was resuspended in 1 mL MiR05 medium (10 mM KH2PO4, 3 mM MgCl2, 500 µM EGTA, 60 mM lactobionic acid, 20 mM taurine, 110 mM sucrose, 1 g/L BSA, and 20 mM HEPES, pH 7.1). Respiratory oxygen flux was assessed at 37°C in MiR05 containing 2 μM digitonin and expressed in picomoles of O2 per second per mg of tissue. L-octanoylcarnitine (1 mM; Sigma), pyruvate (5 mM; Sigma), and malate (2 mM; Sigma), Glutamate (10 mM; Sigma-Aldrich) and succinate (10 mM; Sigma-Aldrich) were added to test complex I and II respiration. Finally, 2 mM GDP (Sigma) was used to estimate the GDP-sensitive respiration.

### Glucose and insulin tolerance test

Glucose tolerance tests were performed following the recommendations of ^44^ in mice after a 6h overnight fast. Glucose was injected intraperitoneal (1.5 g/kg BW), and blood glucose concentrations were measured after 0, 15, 30, 60, 90, and 120 min using a glucometer. Insulin tolerance tests were performed after 2h of fasting. After determination of basal blood glucose concentrations, each mouse received an intraperitoneal injection of insulin (0.75 U/kg BW), and blood glucose concentrations were measured using Optium Neo (Abbott, Ireland) after 15, 30, 60, 90, 120, and 150 min.

### Assessment of mitochondrial morphology via Transmission Electron-Microscopy (TEM)

Freshly excised brown adipose tissues were finely minced and subjected to primary fixation in a solution containing 1% glutaraldehyde, 2.5% paraformaldehyde, 100 mM cacodylate buffer (pH 7.4), 6 mM CaCl₂, and 4.8% sucrose at 4 °C overnight. Following three washes with 100 mM cacodylate buffer (pH 7.4), samples were immersed in a secondary fixation medium with cacodylate buffer and 2% osmium tetroxide at room temperature for 1 h. After thorough washing, the tissues were pre-stained with saturated uranyl acetate for 50 min at room temperature, dehydrated through a graded acetone series, embedded in Epon epoxy resin, and polymerized at 60 °C for 48 h. Ultrathin sections were examined using a JEM-1400-FLASH Transmission Electron Microscope (JEOL JEM-1400, JEOL, Ltd., Tokyo, Japan). Mitochondrial ultrastructure was analyzed using ImageJ software. The maximum length distribution was estimated for each group using the Seaborn library^75^.

### Chromatin immunoprecipitation (Chip) assays

Chromatin immunoprecipitation was performed according to the manufacturer’s protocol (available at www.abcam.com/protocols/immunoprecipitation-protocol), with some modifications. Briefly, adipocytes were incubated in cross-linking solution (1% formaldehyde) at room temperature for 10 minutes. The cells were then washed twice with 1x PBS and detached from the flask. Subsequently, the cells were resuspended in SDS lysis buffer with protease inhibitors and incubated on ice for 10 min. Chromatin fragmentation was performed by sonication in sodium dodecyl sulfate (SDS) lysis buffer using a Sonic VibraCell. Sonication was performed to obtain DNA fragments with an average size of 150–800 bp. Immunoprecipitation was performed using an anti-E4BP4 antibody (Cell Signaling, mAb #14312). IgG was used as a control. The precipitated DNA was analyzed by qPCR using primers (Table S4) targeting the regulatory region of Cers6 (enhancer).

### Site-directed mutagenesis

Site-directed mutagenesis was performed using a Q5 Site-Directed Mutagenesis Kit (NEB) to introduce the desired mutation into the target gene (E4bp4).

Mutagenic primers were designed to incorporate mutations at the center of the primers (Table S4). Point mutations were introduced via site-directed mutagenesis to replace asparagine (Asn, AAC) with lysine (Lys, AAG) and glutamic acid (Glu, GAA) with glycine (Gly, GGA), with the aim of evaluating the effects of altered charge and side chain properties on protein function, following a previously published structure analysis^31,32,76^. The reaction mixture included 50-100 ng of plasmid DNA (Addgene cat. no 34572) as the template, 0.5 μM of each mutagenic primer (forward and reverse), Q5 hot start and 2X master mix following the manufactures’ instructions. After amplification, the reaction was treated with Kinase, Ligase and DpnI (provided in the kit) to digest the parental methylated plasmid DNA, leaving only the newly synthesized, mutated plasmid. The digested plasmid was then transformed into NEB 5-alpha via heat-shock transformation.

### LC-MS lipidomic analyses

The lipid extracts stored at –80°C were resuspended in 150 µL of an acetonitrile:isopropanol mixture (70:30 (v:v)) of UPLC grade, followed by agitation. The resuspended samples were centrifuged at 16,000 × g for 5 min, and 100 µL of the supernatant was transferred to 2 mL glass vials. From each analytical sample, a 20 µL aliquot was taken and pooled. These pools were used as quality control samples for instrumental and sample stability (QCs), which were run after every 5th analytical sample in the sample sequence. All the samples and pools were placed in an Acquity iClass UPLC sample manager (Waters) at 4°C. The UPLC system was coupled to a quadrupole time-of-flight (QTOF) system equipped with an ESI source (Xevo G2-XS QTof, Waters). From each lipid sample, 5 µL was injected onto a 100 x 2.1 mm UPLC CSH C18 column packed with 1.7 µm particles (Waters). The UPLC flow rate was set to 200 µL/min, and the buffer system consisted of buffer A (10 mM ammonium formate in acetonitrile:UPLC water (70:30, (v:v))) and buffer B (10 mM ammonium formate in UPLC isopropanol:acetonitrile (90:10, (v:v))). The UPLC gradient was as follows: 0 – 0.80 min 60% A, 0.80-10 min 60 – 0% A, 10-16 min 0% A, 17 min 0 – 60% A, 17-22 min re-equilibrating at 60% A. This resulted in a total run time of 22 minutes per sample. The mass spectrometer was operated in the negative ionization mode within the mass range of m/z 50-2000. The ESI source was operated with a spray voltage of 2.37 kV in negative ionization mode. Ceramide standard, Cer (16:0) (Avanti Polar Lipids, USA) with a molecular weight of 537.512 g/mol, was used as a reference for retention time and m/z identification under the method’s conditions. The ion was identified as the formate adduct (M+HCOO) − with an m/z of 582.51, eluted at a retention time of 9.33 min. An analytical curve for the relative quantification of Cer(d18:1/16:0) in the samples was generated over the concentration range of 12.5 ppm to 50 ppm, with the curve obtained from 3 different concentration points (12.5 ppm, 25 ppm, and 50 ppm), each analyzed in triplicate. The following equation was derived: y = 1.760 × − 20.232, R² = 0.9935. Relative quantification of ceramide was performed on each sample based on the area under the corresponding ion curve, and an estimate of the concentration of the molecule in the sample was obtained using the calibration curve. The LC-MS data obtained were analyzed using Progenesis QI software (Waters), and untargeted and statistical analyses were conducted using the MetaboAnalyst platform.

### Enhancer data analysis

All enhancers related to the regulatory region of Cers6 were downloaded from the database enhanceratlas.org and aligned in the Genome browser UCSC server across different species, including humans and mice. The alignment was generated by the default method using the region established for *the musculus* (65 kb to TSS of Cers6).

### ChIP-seq analysis

ChIP-seq data for PRDM16 and PRDM16 knockout (KO) were reanalyzed from a previously published study^54^ available at the Gene Expression Omnibus (GSE63965). ChIP-seq data for histone markers (H3K27ac and H3k4Me1) in brown adipose tissue were downloaded from encode project (www.encodeproject.org). ChIP-seq reads were aligned using the Bowtie software ^77^. SAM files were then converted into BAM files. Next, peak calling was determined using MACS^78^. Wig to bigWig files were generated, and peaks were visualized using the UCSC Genome Browser. PRDM16 genome-wide binding sites in brown adipose tissue were defined using the Genomic Regions Enrichment of Annotations Tool (GREAT)^79^.

### RNA preparation and determination of mRNA levels

Total RNA was extracted from cells using TRI reagent (Sigma). CDNA will be synthesized using High-capacity cDNA Reverse transcription kit (Thermo Fisher Scientific) according to the provided protocol. qPCR was performed using Sybr Green (Applied Biosystems) or TaqManTM (Thermo Fisher). The primer sequences and probes for the target genes are listed in Table S4.

### RNA preparation for RNA-seq

RNA was extracted from the tissues using the PureLink RNA kit (Invitrogen). A Bioanalyzer 2100 system (Agilent Technologies) was used to examine RNA integrity. The sequencing library was built using the Library Prep Kit for Illumina® and sequencing using the platform NovaSeq PE150.

### Bioinformatics analyses of RNA-seq

The quality control of the sequences was examined using FastQC (available online at: http://www.bioinformatics.babraham.ac.uk/projects/fastqc). Clean data were obtained using Fastp^80^ to remove low-quality reads. The data were then mapped with the *Mus musculus* mm10 genome reference using HISAT2 (v2.1.0)^81^. The read counts for each gene were calculated using iRNA-seq software. Differential gene expression was identified by DESeq2 (version 1.10.1). A heatmap was generated using the pheatmap (R package) to identify differentially expressed genes. The R package ggplot2 (version 3.3.4) was used to generate volcano plots. Finally, PathfindR^82^ was used for enrichment analysis.

### Bioinformatics analyses of IMAGE

Data acquisition and preprocessing. RNA-seq and ATAC-seq data were obtained from NCBI using the public Gene Expression Omnibus database (GSE181443). The reads obtained were subjected to quality control using the Fast QC tool (available online at: http://www.bioinformatics.babraham.ac.uk/projects/fastqc). Low-quality fragments and sequencing adapters were removed using Trimmomatic. Alignment was performed using the Bowtie2 tool, where the reads were aligned using the mouse mm10 reference genome. Standardization of the sequencing data. RNA-seq data aligned against a reference genome were annotated using iRNA-seq^83^. All data processing was carried out in a Bioconda environment using the feature counts package as a basis. ATAC-seq data already aligned against a reference genome were standardized using the Homer package (available at http://biowhat.ucsd.edu/homer) features: makeTagDirectory, findPeaks, mergePeaks, and annotatePeaks). Integration and analysis of the generated data. The files containing RNA-seq and ATAC-seq data were integrated using IMAGE^51^ in the Bioconda environment following the recommendations of the author. A file containing the gene expression in RPKM was used for RNA-seq analysis. The files containing information on binding sites for different transcription factors were submitted to GREAT^79^ for the annotation of genes related to each site.

### Bioinformatics analyses of Single-cell RNA-sequencing (scRNA-seq)

scRNA-seq data available in^84^ were re-analyzed using Seurat. Low-quality cells and lowly expressed genes were excluded. Data were normalized using the LogNormalize method and highly variable genes were identified using the VST method. A gene expression matrix was generated for downstream dimensionality reduction, clustering, and differential expression analyses.

### Data analysis and graphs

Experiments on the cultured cells were conducted using at least three independent replicates. Comparisons between two independent groups were performed using an unpaired two-tailed Student’s t-test. A paired two-tailed Student’s t-test was used for comparisons between two groups with matched sample pairs. When comparing datasets with more than two groups, one-way analysis of variance (ANOVA) followed by Tukey’s post-hoc multiple comparison test was employed. Statistical analysis for differential expression was performed using a two-sample t-test with Benjamini-Hochberg (BH, FDR of 0.05) correction for multiple testing. The schematic figures were obtained using Biorender.com.

