## Supplementary material for "E4BP4 Safeguards Brown Fat Mitochondria from Obesity-Induced Fragmentation via Ceramide Repression": reviewers' comments

### Revision Plan

Review COMMONS

**Manuscript number:** RC-2025-03113
**Corresponding author(s):** Carlos, Sponton

*[The "revision plan" should delineate the revisions that authors intend to carry out in response to the points raised by the referees. It also provides the authors with the opportunity to explain their view of the paper and of the referee reports.*

*The document is important for the editors of affiliate journals when they make a first decision on the transferred manuscript. It will also be useful to readers of the reprint and help them to obtain a balanced view of the paper.*

*If you wish to submit a full revision, please use our "[Full Revision](Full Revision)" template.* ***It is important to use the appropriate template to clearly inform the editors of your intentions.****]*

#### 1. General Statements [optional]

*This section is optional. Insert here any general statements you wish to make about the goal of the study or about the reviews.*

We appreciate the reviewers' assessment of the significance of our work and would like to highlight where we believe the novelty of this study lies. Our findings identify E4BP4 as a key transcription factor that maintains mitochondrial homeostasis by restraining the overactivation of biological pathways – such as de novo ceramide synthesis – that are known to drive mitochondrial oxidative dysfunction in the context of obesity. We fully acknowledge that the link between C16:0 ceramide and mitochondrial fragmentation has been previously established. However, to our knowledge, our study is the first to connect this phenomenon to a transcriptional safeguard mechanism, thereby providing a new layer of understanding of how transcription factors preserve mitochondrial integrity and function in brown adipocytes. We believe this conceptual advance adds significant value to the field by framing E4BP4 as a transcriptional "guardian" of mitochondrial homeostasis.

#### 2. Description of the planned revisions

*Insert here a point-by-point reply that explains what revisions, additional experimentations and analyses are planned to address the points raised by the referees.*

**Reviewer #1 comment:**
Figures B: Sample size of EE experiments is too low to draw any meaningful conclusions or to know for certain if the data are reproducible. Small sample sizes, likely coming from one litter and one batch of AAV are prone to type I error.

**Response:** We agree with reviewer observation that increasing sample size is essential to confirm reproducibility and robustness. We have therefore planned to repeat the EE experiments with a larger number of mice per group, derived from independent litters and AAV preparations, in order to strengthen the statistical power and validate the phenotype observed in the current study.

**Reviewer #1 comment:**
Figure 3I: Why do cells (none of the groups) show no response to NE stimulation? Please clarify or provide potential mechanistic insight. Perhaps the cells were not differentiated well.

**Response:** We agree that the absence of a robust NE response in Figure 3I requires further clarification. To address this, we have planned to repeat the in vitro oxygen consumption assay to confirm the phenotype presented in the study.

**Reviewer #1 comment:** Figures 3I vs 5N. There is a striking discrepancy between these panels. In both, cells were treated with palmitate for 6 h, yet the NE and CCCP responses differ significantly. Are these the same cell types and conditions? Please reconcile the differences.

**Response:** We would like to clarify that Figures 3I and 5N represent different experimental systems: Figure 3I shows data from primary brown adipocytes with E4bp4 transgene overexpression, whereas Figure 5N shows data from immortalized brown adipocytes with Cas9-mediated mutation of a 65 kb Cers6 enhancer site. Given the distinct cell types and genetic manipulations, a direct comparison between these two panels is not appropriate. Nevertheless, we agree that confirming the consistency of the phenotype across systems is important. To address this, we have planned to repeat oxygen consumption assays in both models to further validate the reproducibility of the observed effects.

---

**Reviewer #2 comment:** A key experiment is missing: does adding C16:0 block the mitochondrial benefits of E4BP4-OE?

**Response:** We thank the reviewer for this excellent suggestion. We agree that a rescue experiment is important to directly test whether C16:0 affects the mitochondrial benefits of E4BP4. To address this, we have planned to perform a co-overexpression of E4bp4 and Cers6 in brown adipocytes. The readouts will include mitochondrial morphology and oxygen consumption, enabling us to determine whether restoration of C16:0 production mitigates the protective mitochondria effects of E4BP4 overexpression. This experiment will provide direct mechanistic confirmation of the proposed model.

**Reviewer #2 comment:** Whether PRDM16-OE mimics the effects of E4BP4 to induce p-Drp1 is not shown.

**Response:** We thank the reviewer for this valuable suggestion. We agree that testing whether PRDM16 overexpression mimics the effects of E4BP4 on p-Drp1 is important to strengthen the mechanistic link between these transcription factors in terms of regulation of mitochondrial fragmentation. To address this, we have planned to include a Western blot analysis of p-Drp1 in the PRDM16-OE in brown adipocytes.

#### 3. Description of the revisions that have already been incorporated in the transferred manuscript

*Please insert a point-by-point reply describing the revisions that were already carried out and included in the transferred manuscript. If no revisions have been carried out yet, please leave this section empty.*

**Reviewer #1 comment:**
Figure 1F: There is an unexpected dip in gene expression at cold exposure days 3 and 7, followed by a rebound at day 14. Is this fluctuation biologically meaningful or technical?

**Response:** We thank the reviewer for this thoughtful observation. A previous study demonstrated that E4bp4 (Nfil3) expression displays an early increase (at 2 hours), followed by a decrease in magnitude – while still remaining significantly higher than control – during beige adipocyte differentiation in response to forskolin treatment (DOI: 10.1016/j.molmet.2022.101619). The authors of that study suggested that E4bp4 may contribute to a second wave of cAMP-driven beige adipocyte differentiation. However, in the context of our work, further discussion on whether the fluctuations in BAT E4bp4 expression observed during cold exposure reflect biological regulation would be speculative. Importantly, despite these oscillations across time points, E4bp4 expression remained statistically significant compared with control, supporting the robustness of our findings. We have now introduced this observation in the Results section of the revised manuscript.

**Reviewer #1 comment:** Figures 2H and 2I (GTT): How was the AUC calculated? The GTT and ITT curves appear largely parallel aside from fasting differences. If total AUC was used instead of incremental AUC, it may overstate group differences. The recommended method is outlined in [DOI: 10.1038/s42255-021-00414-7]. Also, since insulin's half-life is ~10 minutes, later differences in the ITT curve likely reflect counterregulatory responses driven by hepatic gluconeogenesis.

**Response:** We would like to clarify that in our original manuscript we had already calculated the area of the curve (AOC) rather than the area under the curve (AUC), following the recommended approach (DOI: 10.1038/s42255-021-00414-7). Specifically, the AOC was derived by subtracting the baseline glucose value from each subsequent time point, ensuring that the analysis reflects incremental changes rather than absolute glucose levels. We have now made this description more explicit in the revised version to avoid any ambiguity.

**Reviewer #1 comment:** Figure 4F: How was mitochondrial fragmentation quantified? Please ensure that the ROI boxes shown in zoomed panels match the same region in size and shape - this applies throughout the manuscript.

**Response:** We thank the reviewer for this valuable comment. To improve the quality and interpretation of the data, we have now included a quantitative analysis of mitochondrial morphology parameters associated with Figure 4F **(Figure S4B)**. Specifically, we analyzed:

- **Mitochondrial volume (µm³):** reflecting overall mitochondrial size.
- **Surface area (µm²):** reflecting membrane expansion.
- **Sphericity index:** indicating morphological rounding, which increases with fragmentation.
- **Number of branches and branch junctions per mitochondrion:** reflecting mitochondrial networking and fusion.

Related to Figure 4F

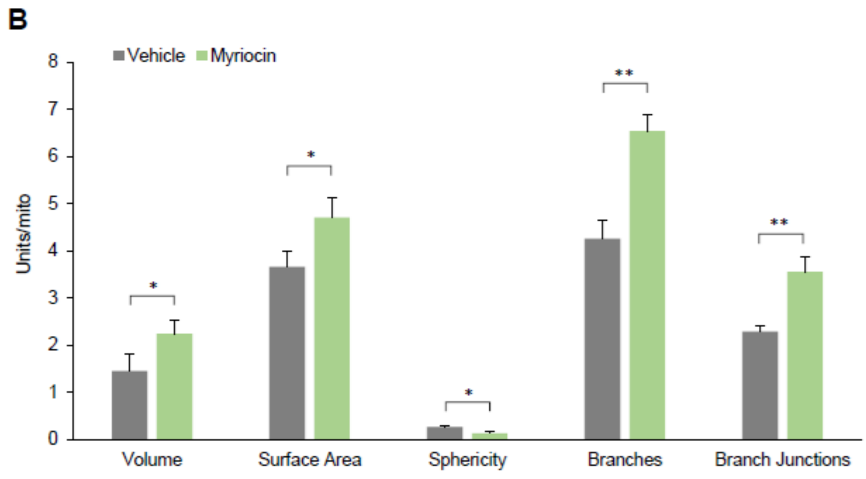


Myriocin treatment preserved mitochondrial volume and surface area, reduced sphericity, and increased both the number of branches and branch junctions, reflecting maintenance of a more interconnected mitochondrial network.

Additionally, we verified that the ROI boxes shown in the zoomed panels are consistent in both size and shape across groups, as requested. We have now introduced this observation in the Methods section of the revised manuscript.

**Reviewer #1 comment:** Figure 3A: The claim that one group contains smaller mitochondria is not convincing. Both small and elongated mitochondria appear in each group. Moreover, it is unclear whether these minor differences are of any physiological relevance or whether they drive phenotypes.

**Response:** We respectfully disagree with this observation and would like to clarify a few points.

1. We have already demonstrated a statistically significant difference in mitochondrial length between E4bp4-OE and control groups (Figures 3B and 3C). This was based on a random, unbiased analysis, which consistently confirmed longer mitochondria in E4bp4-OE compared with control.

2. Some degree of variability in mitochondrial length is expected in electron microscopy analyses, particularly because mitochondria from multiple cell types within iBAT are captured. It is important to note that the protective action of E4bp4 against mitochondrial fragmentation occurs specifically in brown adipocytes, where the transgene is expressed under the control of the adiponectin promoter.

3. To address the potential confounding heterogeneity of iBAT mitochondria, we performed complementary cell-autonomous analyses in vitro, allowing us to directly compare mitochondrial dynamics in E4bp4-OE versus control brown adipocytes. This analysis further confirmed that E4bp4-OE prevents lipid overload – induced mitochondrial fragmentation in brown adipocytes.

4. Finally, we emphasize that several studies have demonstrated that changes in mitochondrial dynamics, particularly under high-fat diet conditions, disrupt systemic energy homeostasis (DOI: 10.1016/j.cmet.2017.05.010; DOI: 10.1016/j.cell.2019.05.008; DOI: 10.1038/s42255-024-00978-0). Therefore, the differences we report are biologically meaningful in the broader context of mitochondrial dynamics and metabolic disease.

**Reviewer #1 comment:** Figure 3E: The claim that confocal microscopy reveals palmitate-induced mitochondrial fragmentation is difficult to discern. The images lack clear morphological differences.

**Response:** We thank the reviewer for this observation. To improve the interpretation of these results, we have now included a quantitative analysis of mitochondrial morphology parameters associated with **Figure 3E.** Specifically, we measured:

- **Mitochondrial volume (μm$^3$):** reflecting overall mitochondrial size.
- **Surface area (μm$^2$):** reflecting membrane expansion.
- **Sphericity index:** indicating morphological rounding, which increases with fragmentation.
- **Number of branches and branch junctions per mitochondrion:** reflecting mitochondrial networking and fusion.

As shown in the new analysis **(Figure S4A)**, palmitate treatment reduced mitochondrial volume, surface area, branches, and branch junctions, while increasing sphericity, consistent with a more fragmented phenotype in control cells. In contrast, these effects were significantly attenuated in E4bp4-OE cells, supporting our conclusion that E4BP4 overexpression protects against lipid overload-induced mitochondrial fragmentation. This text was added in the Results section of the revised manuscript.

Related to Figure 3E

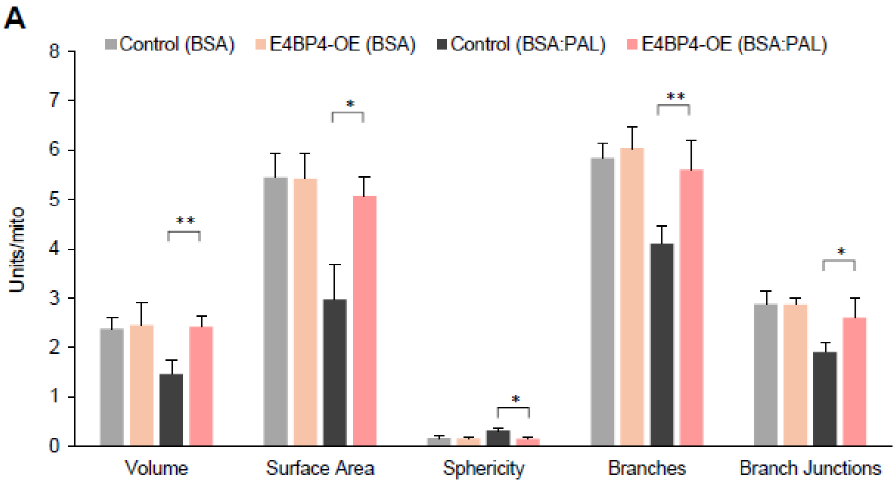


We believe this additional analysis strengthens the robustness of our findings and provides clear quantitative evidence for the morphological changes that were less apparent from qualitative image inspection alone.

**Reviewer #1 comment:** Figure 3G: Dendra2-labeled mitochondria appear unaffected by palmitate, raising concern about the robustness of the effect across readouts.

**Response:** We respectfully disagree with this observation. As shown in **Figure 3G** (bar graphs), palmitate-treated brown adipocytes exhibited a clear reduction in mitochondrial co-localization, which reflects lower levels of fused mitochondria, in the control group compared with E4bp4-OE. Importantly, no difference in mitochondrial co-localization was observed between the two groups under vehicle-treated conditions. This indicates that E4bp4 overexpression does not promote mitochondrial fusion per se, but rather prevents lipid overload – induced mitochondrial fragmentation. We also note that the representative images presented in **Figure 3G** are single snapshots taken from a time-lapse assay of mitochondrial dynamics. To further illustrate this effect, we direct the reviewer to the supplementary video accompanying this experiment, which clearly demonstrates the differences in mitochondrial behavior over time.

**Reviewer #1 comment:** Figure 5H: Were E4BP4 expression levels equivalent between WT and mutant cells? Quantification should be shown. Figure 5H: Were E4BP4 expression levels equivalent between WT and mutant cells? Quantification should be shown.

**Response:** We thank the reviewer for this important point. We have now added the quantification of E4bp4 mRNA levels in cells transduced with either the non-mutated vector (control) and the vector carrying a mutation in the E4bp4 DNA-binding domain **(Figure S5)**. The data show no significant difference in E4bp4 expression between the two groups.

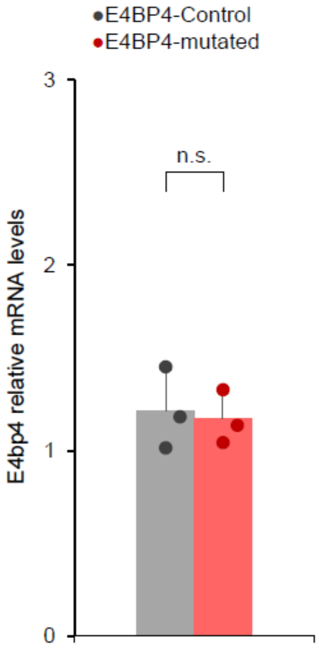


---

**Reviewer #2 comment:** The evidence of mitochondrial fragmentation is not convincing. In the reviewer's opinion, Figures 3E, 3G, 4F, and 5M demonstrated a decrease in mitochondrial quantity, but not fragmentation.

**Response:** We thank the reviewer for this observation. We have already addressed the comments from reviewer #1 (above) regarding Figures 3E, 3G and 4F related to measurements of mitochondria fragmentation. To strengthen the interpretation of these results, we have also performed a quantitative analysis of mitochondrial morphology parameters associated with **Figure 5M**. Specifically, we measured:

- **Mitochondrial volume (μm³):** reflecting overall mitochondrial size.
- **Surface area (μm²):** reflecting membrane expansion.
- **Sphericity index:** indicating morphological rounding, which increases with fragmentation.
- **Number of branches and branch junctions per mitochondrion:** reflecting mitochondrial networking and fusion.

As shown in the new analysis **(Figure S4C)**, palmitate treatment significantly reduced mitochondrial volume, surface area, and branching, while increasing sphericity, consistent with enhanced mitochondrial fragmentation in control cells. Notably, these changes were significantly blunted in the Cers6 enhancer edited cells (EΔ), supporting our conclusion that disruption of Cers6 protects against lipid overload–induced mitochondrial fragmentation**.** This text was added in the Results section of the revised manuscript.

Related to Figure 5M

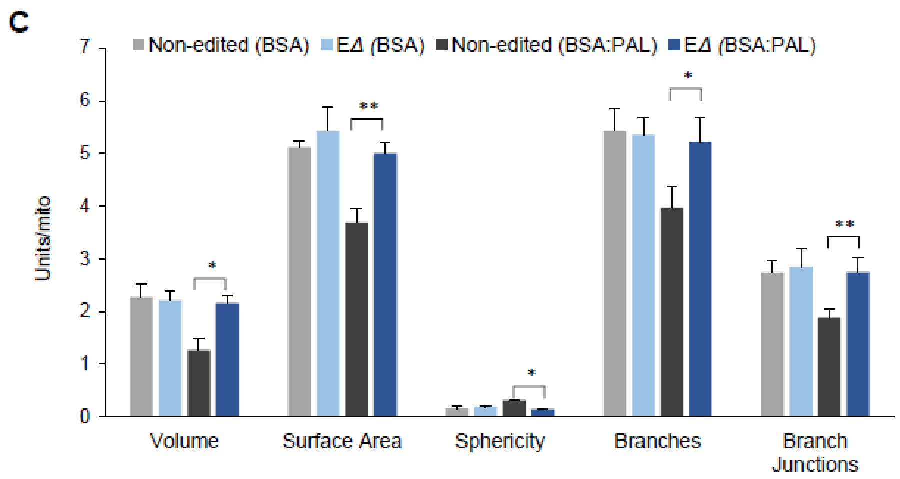


Regarding the reviewer's understanding of a "decrease in mitochondrial quantity, but not fragmentation," we respectfully disagree. The analyses performed for Figures 3E, 3G, 4F, and 5M clearly demonstrate that E4bp4 overexpression (E4bp4-OE) prevents lipid overload –induced mitochondrial fragmentation.

In relation to mitochondrial quantity, our data do not support differences in mitochondrial biogenesis between groups. Specifically, the expression of thermogenic and mitochondrial biogenesis genes **(Figure S2G)** as well as the mitochondrial-to-nuclear DNA ratio **(Figure S3D)** showed no significant changes, indicating that mitochondrial biogenesis is not altered.

Alternatively, it is possible that E4bp4 prevents mitophagy, as our results **(Figure 3H)** show that E4bp4-OE protects against lipid overload-induced mitochondrial depolarization. In this regard, previous studies have demonstrated that fragmented and depolarized mitochondria are targeted for degradation through mitophagy (DOI: 10.2337/db07-1781; DOI: 10.1074/jbc.M111.242412). While this explanation is consistent with our findings, we acknowledge that it remains speculative at this stage and, although interesting, is beyond the scope of the current study.

**Reviewer #2 comment:** It is confusing whether the association shown in Figure 1C is a positive or an inverse association.

**Response:** We thank the reviewer for pointing out this source of confusion. **Figure 1C** represents common variant associations for E4BP4, where the y-axis indicates the strength of association (-log10 p-value) rather than the direction (positive or inverse) of the effect. We have clarified this in the revised manuscript to avoid misinterpretation. The associations indicate that genetic variants in E4bp4 are positively linked with anthropometric traits such as weight, BMI, and waist–hip ratio.

#### 5. Description of analyses that authors prefer not to carry out

*Please include a point-by-point response explaining why some of the requested data or additional analyses might not be necessary or cannot be provided within the scope of a revision. This can be due to time or resource limitations or in case of disagreement about the necessity of such additional data given the scope of the study. Please leave empty if not applicable.*

**Reviewer #2 comment:**
It would be worthwhile to investigate whether in vivo knockdown of E4BP4 blunts the Cers6-suppressing effects of PRDM16-OE.

**Response:** We agree that assessing in vivo loss-of-function of E4bp4 in the context of Prdm16 overexpression would be highly informative. At present, this experiment is technically not feasible, as it would require the generation and characterization of complex *in vivo* models beyond the scope of the current study. Nevertheless, we are actively considering this as a future direction. In the meantime, we believe that the *in vitro* experiments in brown adipocytes provided here are sufficient to establish the mechanistic relationship between E4BP4 and PRDM16 in the regulation of Cers6 expression.

**Reviewer #2 comment:** Whether E4BP4-OE affects cold tolerance in mice is now shown.

**Response:** We thank the reviewer for this thoughtful comment. In our study, we performed an iBAT-specific E4bp4 gain-of-function assay because we observed a downregulation of E4bp4 expression in the context of obesity. The rationale for this approach was to rescue E4bp4 expression in iBAT and thereby evaluate its systemic and mechanistic effects under obesogenic conditions. We recognize that a gain-of-function assay during cold challenge would further enhance E4bp4 expression and, while interesting, this would more directly address the role of E4bp4 in thermogenic regulation rather than in obesity-related metabolic dysfunction. For this reason, we believe that a detailed investigation of E4bp4 in cold-induced thermogenesis is an important but separate question that lies beyond the scope of the current study.
